# Explainable Decoding of Sensorimotor Communication in Joint Object Manipulation

**DOI:** 10.64898/2026.08.17.745075

**Authors:** Yiming Liu, Dorian Verdel, Raz Leib, Etienne Burdet, David W. Franklin

## Abstract

Humans often collaborate under asymmetric information, for example when two people carry a table and only one knows the destination. They coordinate without speech using cues from movement kinematics, interaction forces, and object states. Characterizing this sensorimotor communication is difficult because these signals both execute the task and convey information, whose meaning is context-dependent. Here, we investigated a virtual table-carrying task where one partner knew the target while the other inferred it from visuo-haptic feedback. Participants flexibly adapted kinematic and haptic cues across contexts to convey intention. We introduce an explainable machine-learning framework that decodes intent from ongoing multimodal signals and quantifies where individual features are informative. Incorporating the decoded signals into a drift-diffusion model accurately predicted the uninformed partner’s target choices and decision times. Together, our framework explains how humans communicate through action and offers principles for collaborative robots to infer and express intent through physical interaction.

---

Humans excel at physical collaboration, coordinating seamlessly by continuously predicting and adapting to their partners’ actions [1, 2, 3, 4, 5, 6, 7, 8], while forming internal representations of their partners’ behavior [9, 10, 11]. Achieving fine coordination requires partners to learn each other’s movement intentions, which becomes especially challenging under *asymmetric information*, when one partner lacks full knowledge of the environment (e.g., the goal or an obstacle) and must infer it from their partner’s behavior. Despite this information gap, partners can often coordinate effectively without words, communicating intentions through action by modulating movements and forces, a process referred to as sensorimotor communication [12]. Effective communication may be facilitated by legible behavioral signals that enable observers to infer intent quickly and confidently [13]. Although central to human physical collaboration, the mechanisms by which movement and force jointly convey intent remain largely unknown.

Sensorimotor communication emerges intuitively in humans [14], yet its underlying structure is difficult to characterize. One reason is its inherently multimodal nature: it relies on high-dimensional kinematic and haptic signals [15] such as movement path, speed and force magnitude. These signals are physically coupled and often covary, yet they remain dissociable and can diverge, for example during collisions. Consequently, changes in one variable may reflect intentional signaling, mechanical task constraints, or indirect consequences of modulating another variable. Moreover, actions in physical collaboration simultaneously serve the task goal and convey intentions and task-relevant information. Communicative signals are embedded within task execution and must respect the behavioral constraints imposed by the task. Together, these properties make human sensorimotor communication difficult to interpret.

Previous studies show that humans enhance legibility by exaggerating or disambiguating movements relative to alternative goals. For example, they modulate movement trajectories [16, 17, 18, 19], velocity profiles [20, 21], grip aperture [17], movement duration [22], and movement variability [23, 24] (see [12] for a review). Communicative signals are often identified as deviating from energetically optimal movements observed when signaling is unnecessary [16, 25]. Although this line of work is motivated by direct physical collaboration (e.g., transporting a table [26, 27, 28]), most studies have examined visually observable cues in tasks without shared object dynamics or direct physical contact. This constrains the multimodal expression of sensorimotor communication and overlooks the contributions of haptic signals and their interaction with visual cues.

Haptic feedback is a critical communication channel during physical collaboration [29, 30]. Humans use haptic cues to infer a partner’s movement intentions [10, 31], which can improve interpersonal coordination [32, 33] and task performance [34, 35]. These effects depend on factors such as the coupling stiffness [34], individual performance [36], individual motor costs [37], and role distribution within the dyad [32, 33, 38]. In prior studies, haptic feedback arose naturally from physical coupling during task execution, but the tasks did not require participants to actively shape haptic signals to enhance legibility because both partners had full visibility and faced no environmental constraints. Moreover, it remains unclear how haptic signals interact with visual cues during active communication.

Here we examine how intention is encoded in and decoded from multimodal sensorimotor signals during physical collaboration. In complex collaboration, partners may embed intention in force, movement, and object-state signals, creating a high-dimensional signaling space that is potentially redundant, as the same intent can be expressed through different combinations of signals. Meanwhile, the informativeness of each signal may depend on the task structure, local environment, and concurrent signals. We ask three questions. First, how is intent embedded across multimodal sensorimotor signals as a function of contextual factors? Second, does intent expression show consistent patterns within individuals, and are these patterns shared across participants? Third, how can we decode and interpret the information conveyed through sensorimotor communication?

To study multimodal sensorimotor communication, we used a joint transport task in which pairs of participants moved a virtual table freely in a plane toward one of four possible targets while receiving visual and haptic feedback. This setup created a rich set of cues through which partners could communicate flexibly, without being constrained to a predefined signaling strategy. By manipulating whether target information was shared or asymmetric between partners, we examined how humans shaped their behavior to make the target legible.

Interpreting these high-dimensional sensorimotor interactions requires new data processing pipelines (Fig. 1). We used explainable machine learning methods to decode the intended target from ongoing behavior and to quantify the contribution of each variable to intent estimation. Based on the decoded signals, we further modeled how the uninformed participant accumulated evidence over time until they identified the target. This approach provides a quantitative method for studying how intent is communicated through physical interaction.

**Figure 1:**
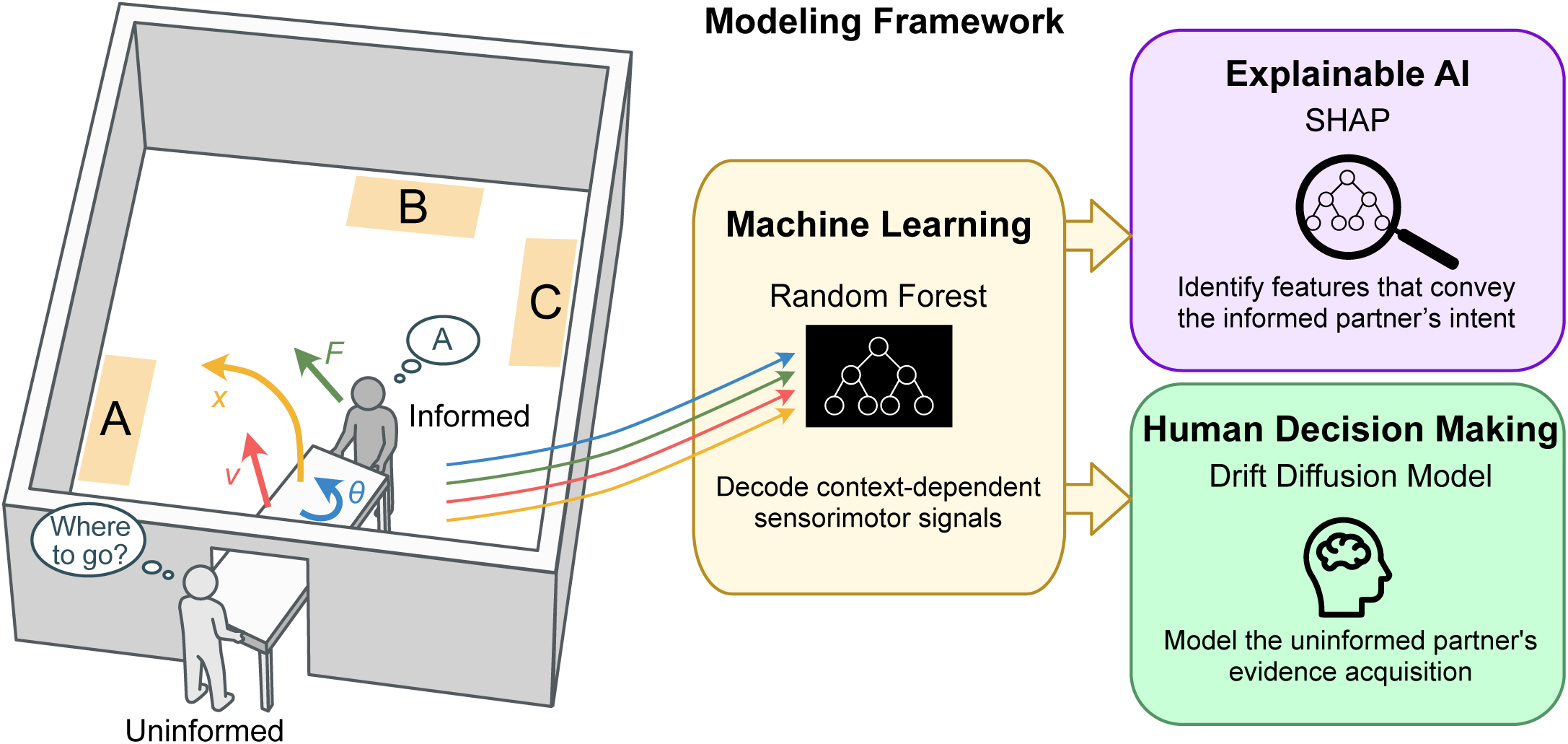
Conceptual overview of the modeling framework. In the asymmetric table-carrying task, only the informed partner knows the target location, while the uninformed partner must infer it from multimodal sensorimotor signals, including object path, velocity, orientation, and interaction force. A random forest model was trained to decode these sensorimotor signals and predict the informed partner’s intended target. We then applied SHAP analysis to explain how each sensorimotor feature contributes to the random forest model’s predictions. Finally, we combined the random forest model with a drift-diffusion model to characterize the uninformed partner’s evidence accumulation and decision-making process.

## 1 Results

Two participants jointly controlled the movements of a table in a four-room virtual apartment using haptic devices, where each partner held an extremity of the table. Each participant viewed the shared task on their own display, but could not see or talk to their partner directly (Fig. 2A). In each trial, four potential target locations were presented, one true target and three decoys. The dyads needed to transport the table from a fixed start position in the left room, through two passages, to the target as quickly as possible. The targets were distributed in the top-right and bottom-right rooms. Two scenes with different target layouts were presented in a pseudo-randomized order across trials. When a participant was informed, the target was highlighted in green with decoys in orange; when uninformed, all potential target locations appeared identically in orange (Fig. 2A). Three experimental conditions varied the information available to each participant and the incentive for early, legible signaling (Fig. 2B). In all conditions, the dyads were rewarded for faster completion times. In the *Symmetric* condition, both participants were informed of the location of the target. In the *Asymmetric* and *Button Press* conditions, only one participant was informed of the location of the target, which they needed to convey to the other uninformed participant through sensorimotor signals. In the *Button Press* condition, the uninformed participant was instructed to press a button as soon as they believed they had identified the target. After the button press, one of two trial types occurred: (i) *Button Press–Transport*, where the dyad continued moving the table to the target, and (ii) *Button Press–Select*, where the table was frozen and the uninformed participant selected the target they believed to be correct. The two trial types were pseudo-randomized within each block and revealed only after the button press. This design encouraged movements that were informative early and compatible with subsequent transport (see Methods, Experimental conditions). In Button Press–Select trials, the trial ended with target selection, and dyads were rewarded based on the time of the button presses for correct selections. In all other trials, the trial ended when the table reached the target, and dyads were rewarded for the actual completion time. General trends of trial-by-trial learning are shown in Supplementary Note 1.

**Figure 2:**
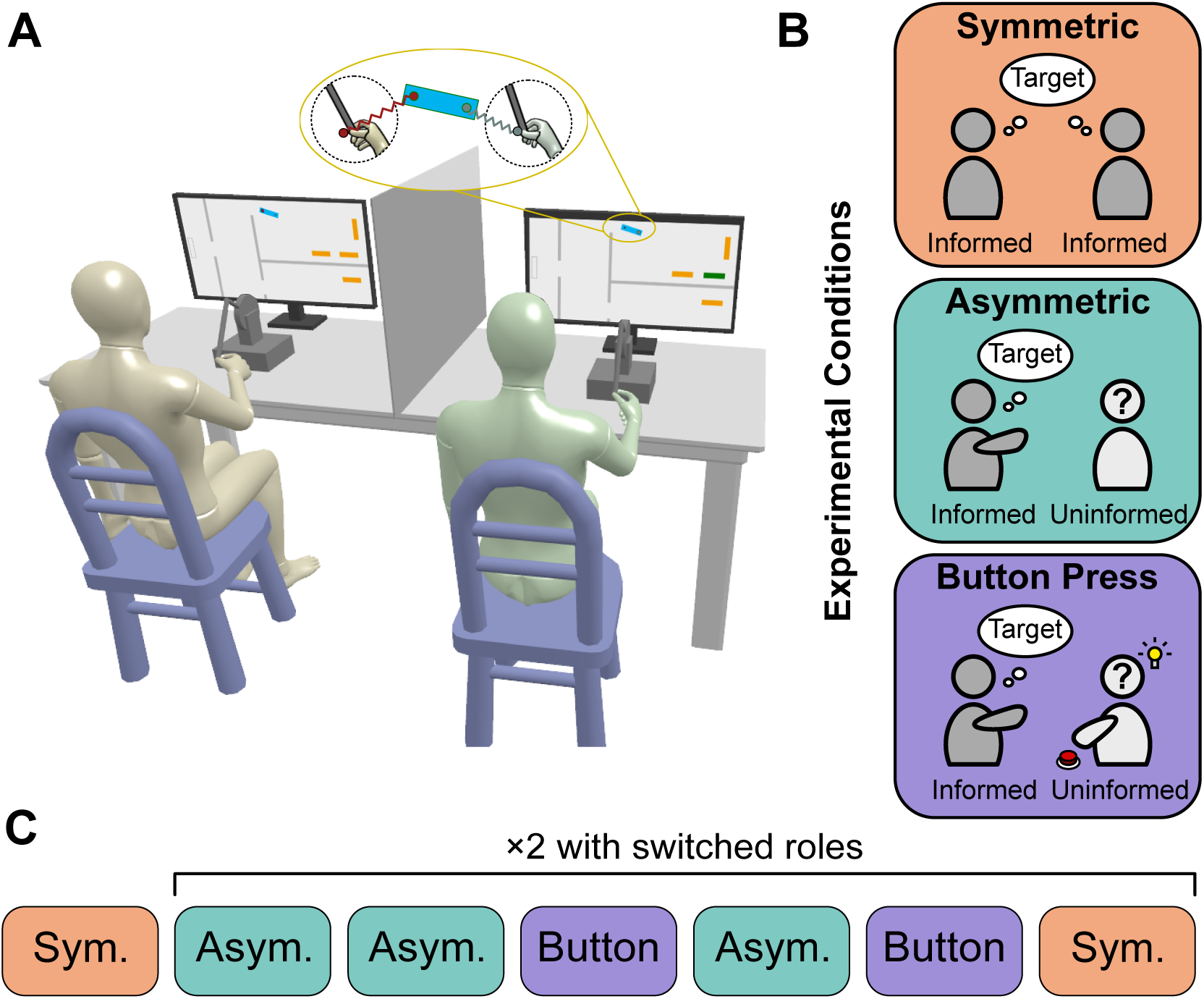
Experimental setup, conditions and protocol. **A.** Experimental setup: two participants sat side-by-side, separated by a curtain, each interacting with a haptic device. The haptic devices were connected to the two ends of a virtual table (blue rectangle) via virtual springs. The virtual apartment was divided into four rooms by virtual walls. Four potential target locations were in the two rooms on the right, with one true target and three decoys. Dyads were instructed to move the table from the start position in the left room to the target, while avoiding collisions with the walls and entering decoy locations. When informed (here on the right), a participant saw the target in green and decoys in orange; when uninformed (here on the left), a participant saw all potential targets in orange. **B.** Experimental conditions: *Symmetric*, both participants were informed of the target location; *Asymmetric* and *Button Press*, one participant was informed and the other was uninformed of the target. In the *Button Press* condition, the uninformed participant pressed a button as soon as they believed they had identified the target. **C.** Experimental Protocol: each dyad performed 13 blocks in total, with each block consisting of 40 co-manipulation trials. The order of block conditions was the same for all dyads. In the *Asymmetric* and *Button Press* conditions, the informed and uninformed roles were swapped between partners halfway through the experiment.

### 1.1 Task performance and effort sharing

To quantify how task conditions shaped performance and dyadic effort, we compared the completion time and total mechanical work performed by the dyad in the final block of each condition (Blocks 5–7,11–13). A repeated-measures ANOVA revealed a significant main effect of condition on completion time (*F*_1.16,18.56_ = 4.28, *p* = 0.048, *η*^2^ = 0.21; Greenhouse-Geisser corrected; Fig. 3A). While the *Symmetric* condition appeared slightly faster, no pairwise comparisons reached statistical significance (all *p >* 0.10).

**Figure 3:**
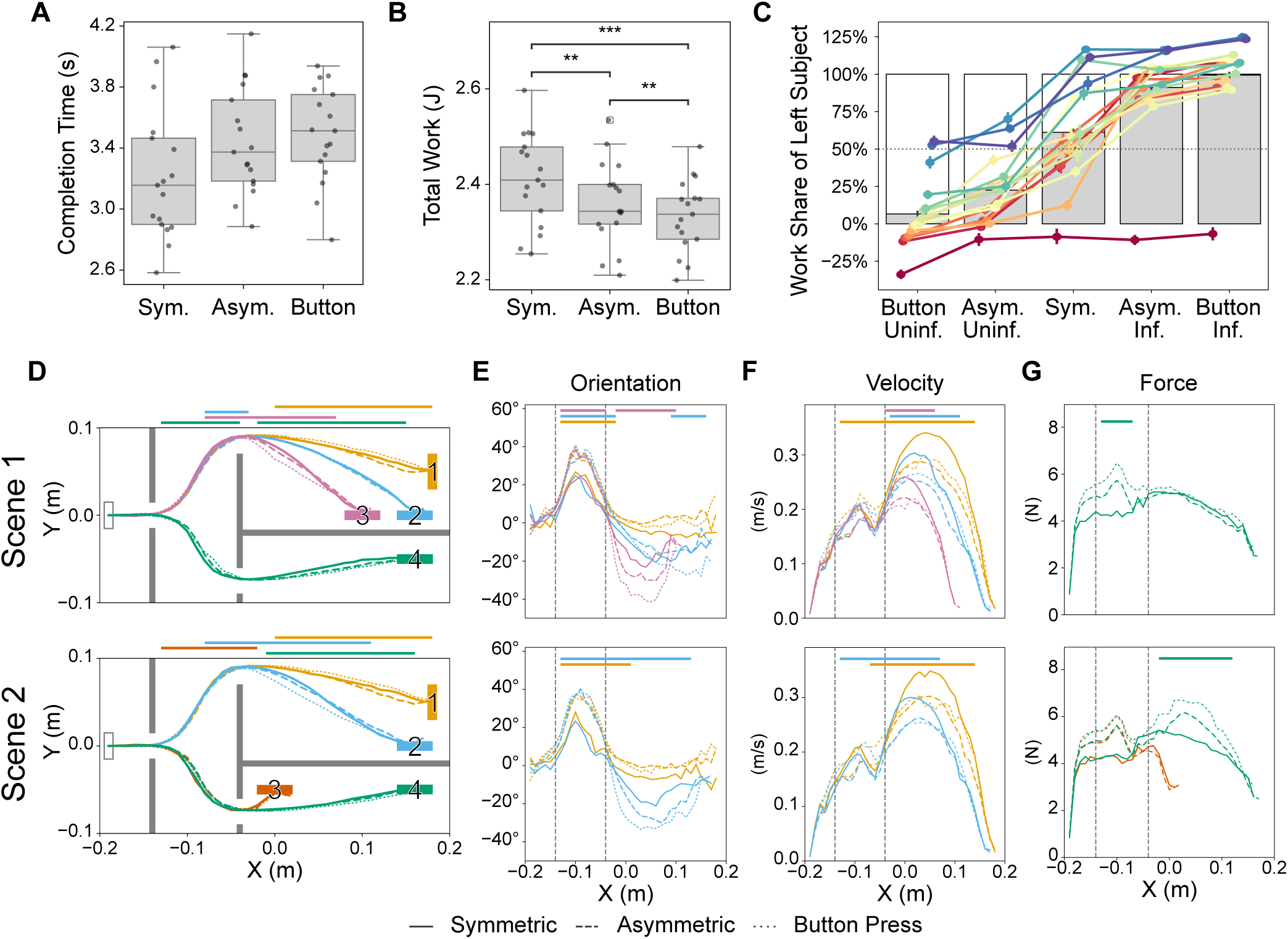
Leader-follower behavior and signaling cues under asymmetric information. **A.** Completion time tended to increase under asymmetric information. **B.** The total mechanical work expended by the dyads decreased under asymmetric information. Asterisks indicate significant pairwise differences. In (A–B), values were calculated from the final blocks of each condition before and after the role swap (Blocks 5–7 and 11–13). Button Press–Select trials were excluded as they did not end at the target. **C.** Share of total work contributed by the left-seated participant across task-role conditions, which refers to the combination of the task condition and the participant’s informed/uninformed role. The gray bars indicate the mean across dyads, and the dotted line marks equal contribution. Each colored line corresponds to one dyad. **D.** Mean table trajectories for each target in Scene 1 (top) and Scene 2 (bottom), color-coded by target. **E–G.** Mean profiles of sensorimotor signals plotted as a function of table position along the *x*-axis for each scene: (E) table orientation, (F) table velocity, and (G) leader force. In (D–G), solid, dashed, and dotted lines denote the *Symmetric*, *Asymmetric*, and *Button Press* conditions, respectively. Horizontal bars at the top indicate *x*-ranges with significant differences between conditions according to non-parametric cluster-based permutation tests (see Methods for details), color-coded by target. The dashed vertical lines indicate the positions of the vertical walls with constrained passages; from left to right, they mark the first and second passages. Each panel displays measures exclusively for targets within one room (top-right or bottom-right) of the scene.

We hypothesized that asymmetric information contexts would motivate sensorimotor signaling, which could require an increase in mechanical work compared to the *Symmetric* baseline. Contrary to our hypothesis, we observed a significant decrease in this energetic expenditure (Fig. 3B). A repeated-measures ANOVA revealed a significant main effect of condition on total mechanical work (*F*_1.22,19.52_ = 15.39, *p <* 0.001, *η*^2^ = 0.49; Greenhouse-Geisser corrected). Subsequent pairwise comparisons confirmed that the Symmetric condition required significantly more work than both asymmetric conditions (both: *p <* 0.01, *d >* 0.56). Additionally, the *Asymmetric* condition required more work than the Button Press–Transport condition (*p* = 0.002, *d* = 0.31). Note that the Button Press–Select trials were excluded from this analysis because the table was frozen at the button press. Similar results were obtained when measuring effort through the total force and the table path length (see Supplementary Note 2). In sum, these results show that efficient sensorimotor signaling was achieved without an increase in effort.

Beyond this overall decrease in mechanical work, its sharing between the partners was shaped by information and task requirements (Fig. 3C). Dyads exhibited a consistent monotonic shift in work sharing across task-role conditions, which refers to the combination of the task condition and the participant’s informed/uninformed role. A repeated-measures ANOVA revealed a significant effect of task-role condition (*F*_2.0,32.0_ = 123.15, *p <* 0.0001, *η*^2^ = 0.89, Greenhouse–Geisser corrected). All post hoc pairwise comparisons were significant (all *p <* 0.001). Effect sizes were smaller for comparisons between *Asymmetric* and *Button Press* within the same information role (*d* = 0.28 for informed; *d* = 0.67 for uninformed), while all other pairwise contrasts had *d ≥* 0.93. In the *Symmetric* condition, work sharing was typically more balanced between partners; in *Asymmetric* and *Button Press*, the informed partner contributed more than the uninformed. This imbalance was stronger in *Button Press* than in *Asymmetric*, which may reflect that the informed partner adopted a more dominant role to generate clearer sensorimotor cues, while the uninformed partner prioritized target inference over contributing to transport. Accordingly, in the rest of this paper, we refer to the informed partner as the leader in *Asymmetric* and *Button Press*, and to the partner with the larger work share as the leader in *Symmetric*.

### 1.2 Multimodal signaling

In this joint manipulation task, partners were mechanically coupled through the object, creating a potentially rich physical channel for communication. In principle, the informed partner could embed target information in multiple variables, such as the table’s path, orientation, or interaction forces. To characterize how behavior changed under asymmetric information, we compared these variables across conditions. The *Symmetric* condition was used as a baseline, as it reflects efficient transport without the need for signaling. Deviations from this baseline in *Asymmetric* and *Button Press* conditions may indicate adjustments to improve legibility.

Dyads collaborated to transport the table in two scenes with different target layouts (Fig. 3D). Asymmetric information induced significant changes in table paths in specific regions, generally increasing separation from paths leading to nearby decoys. Furthermore, deviations were generally larger in the *Button Press* condition than in the *Asymmetric* condition (Fig. 3D). Despite these systematic deviations, paths remained broadly similar across conditions, indicating that the dyads preserved transport efficiency by avoiding large detours. This conservative modulation of the path is consistent with the absence of increased dyadic work under asymmetric information (Fig. 3B), suggesting that cues other than path deviations may be used to convey target information. We then examined differences in table orientation, table velocity, and interaction forces between the three conditions. Table orientation profiles showed significant differences between conditions after the second passage. Deviations from the *Symmetric* baseline emerged in the *Asymmetric* condition and were even more pronounced in the *Button Press* condition (Fig. 3E). Table velocity also differed across conditions (Fig. 3F), with the *Symmetric* condition exhibiting higher speeds than the *Asymmetric* and *Button Press* conditions. Finally, leaders applied higher forces in the *Button Press* and *Asymmetric* conditions, particularly between the two passages (Fig. 3G). Comparing the two scenes further revealed that signaling adjustments depended on local environmental context rather than target location alone. *T* 1, *T* 2, and *T* 4 had identical positions across scenes, whereas *T* 3 differed in location. Thus, the same spatial location could correspond to a different local context with different nearby decoys. For instance, in Scene 1, *T* 2 lay between *T* 1 and *T* 3, leading to minimal deviations from the *Symmetric* baseline as shifts in either direction would have been misleading. By contrast, in Scene 2, *T* 2 had the same location but a different local context, with only one competing decoy (*T* 1) on the upper right, leading to downward shifts in path and orientation. Another example was *T* 4 after the second passage: leader force showed little difference across conditions in Scene 1, since no decoys were present. However, in Scene 2, leader force increased under asymmetric information, possibly to further enhance separation from *T* 3. Conversely, targets with similar local contexts showed similar signaling strategies despite different locations. *T* 3 in Scene 1 and *T* 2 in Scene 2 both had competing decoys only to the upper right, and both showed similar downward adjustments. On the other hand, deviations from the *Symmetric* baseline cannot always be attributed to sensorimotor signaling. As shown in Fig. 3C, information asymmetry shifted more work share toward the leader, which could cause behavioral changes. For example, before the second passage, table orientation was more tilted in the *Asymmetric* and *Button Press* conditions (Fig. 3E), which could result from the leader dominating the maneuver, or from table tilt being used as an informative signal, or both. Similarly, the lower speed observed in the *Asymmetric* and *Button Press* conditions likely arose because one partner did not know the target, preventing the dyad from pushing together consistently, rather than because speed itself served as a signal (Fig. 3F).

These results, together with comparisons of other sensorimotor signals (see Supplementary Note 3), suggest that sensorimotor signaling is multimodal, high-dimensional, and context-dependent. However, behavioral differences under asymmetric information may not always reflect signaling alone, motivating the use of more powerful tools to decode it.

### 1.3 Machine learning decoding of sensorimotor communication

Decoding the target from the instantaneous movement state requires combining multiple cues whose meaning can be context dependent. These cues interact non-linearly, which complicates the interpretation of their combined meaning using classical methods. Therefore, we used a Random Forest (RF) classifier to implement context-dependent decision rules by partitioning the feature space, interpret sensorimotor cues, and output probabilities for each target. The probabilities reflect the model’s confidence in each target as the intended goal, and thus the legibility of the instantaneous sensorimotor signals.

The spatial distribution of the Random Forest model’s confidence for each target in Scene 2 is shown in Figure 4A, with the corresponding results for Scene 1 shown in Supplementary Note 4.1. At the start of trials, confidence was near chance level (*∼* 25%), as the initial trajectories were similar across targets and thus uninformative. Confidence increased as the table moved toward the target, with some regions reaching higher confidence earlier than others. In the top-right room, confidence rose faster for *T* 1 along the scene’s upper boundary and for *T* 2 within the lower part. In the bottom-right room, *T* 3 was positioned near the passage and along the route to *T* 4. Accordingly, *T* 3 had high confidence only within a small area around the target, whereas for *T* 4, confidence sharply increased even before the table passed *T* 3, suggesting clear signaling beyond position cues around *T* 3. Confidence dropped sharply when the table position was more consistent with a decoy, as visible near the top-right corner for *T* 2 and the lower-right corner for *T* 3. Overall, although no ground-truth confidence values were available for direct comparison, the model’s outputs qualitatively aligned with our expectations based on the task geometry.

**Figure 4:**
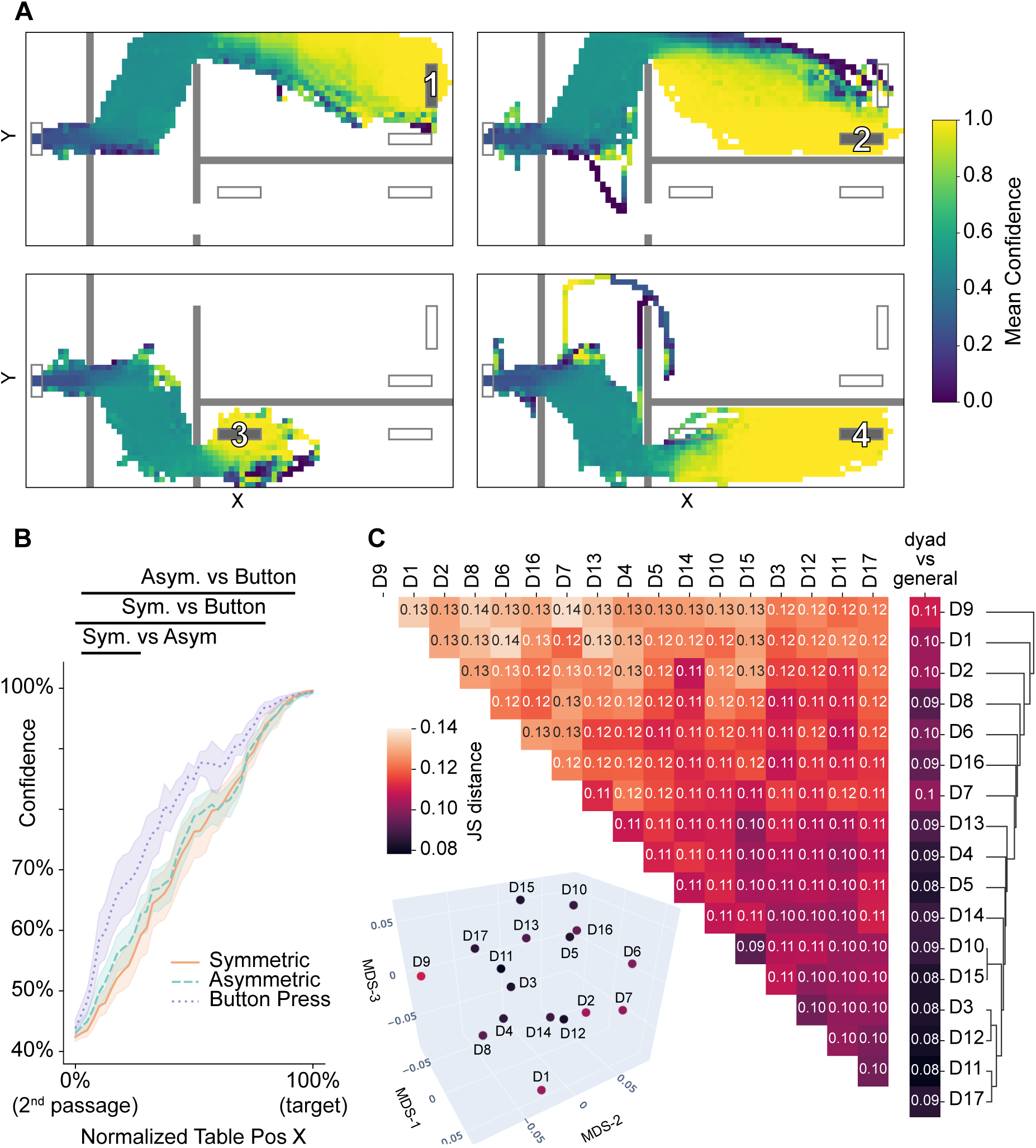
Random Forest decoding of sensorimotor signaling. **A.** Mean confidence of the RF model across the workspace in Scene 2 for each target *T* 1 − *T* 4. The workspace was discretized into 0.5 × 0.5 cm bins, and color indicates the mean predicted probability of the true target within each bin. **B.** Mean model confidence in the true target as a function of normalized table position, from the second passage (0%) to the target (100%), averaged across targets and scenes for each condition. Shaded regions indicate 95% confidence interval across dyads, and horizontal bars at the top indicate significant differences between conditions according to non-parametric cluster-based permutation tests. **C.** Pairwise Jensen–Shannon (JS) distance between dyad-specific RF models, computed from their predictions on a shared dataset. The heatmap shows the mean pairwise JS distance between pairs of dyads, the rightmost column shows the JS distance between each dyad and the general model trained on pooled data. The dendrogram on the right shows agglomerative hierarchical clustering with average linkage, grouping dyads based on the similarity of their model predictions. Inset: 3D metric multidimensional scaling (MDS) embedding of the JS distance matrix, providing a low-dimensional visualization of the distances between dyads.

We next examined how confidence evolved across conditions, averaged across scenes and targets (Fig. 4B). The RF model’s confidence in the true target is plotted as a function of the table’s normalized horizontal position, where 0% corresponds to the second passage and 100% to the target location. Confidence started at similar levels across conditions, with the *Button Press* condition showing slightly higher values. As the table moved toward the target, confidence increased earliest and most strongly in *Button Press*, followed by *Asymmetric* and then *Symmetric*, with significant differences between conditions. This suggests that dyads produced more informative movements when signaling incentives were stronger, with the clearest signaling in *Button Press*, followed by *Asymmetric*, both clearer than the *Symmetric* baseline. In all conditions, confidence approached 100% near the target.

In addition to the general RF model trained on pooled data from all dyads, we also trained dyad-specific RF models on each dyad’s data to assess their individual signaling strategies. The dyad-specific models were evaluated on a shared dataset aggregated from all dyads, and their predictions were compared using the Jensen-Shannon (JS) distance.

The mean pairwise JS distance between each pair of dyads and each dyad compared to the general model is shown in Figure 4C, where smaller JS distances indicate more similar strategies. To characterize how dyads differed in their signaling strategies, we applied agglomerative hierarchical clustering with average linkage based on the JS distance. Dyads that clustered earlier and appeared lower in the dendrogram were more similar to one another, whereas dyads that clustered higher showed more distinct strategies. Moreover, we applied metric multidimensional scaling (MDS) to the JS distance matrix, embedding dyads in a three-dimensional space that approximately preserves their high-dimensional distances (stress=0.124, Fig. 4C inset). Dyads located closer together showed more similar signaling strategies, whereas those farther apart exhibited more distinct strategies. Overall, JS distances remained small and spanned a narrow range (0.08–0.14), indicating that most dyads had similar signaling strategies. A few dyads showed larger distinction to both the general model and other dyads. In particular, D9, D1, and D2 exhibited the largest JS distance, appeared at the top of the dendrogram, and also appeared at the edge of the MDS map. This indicates comparatively distinct strategies. Conversely, dyads near the bottom of the dendrogram (e.g., D10, D15, D3, and D12) were closer to the general model and to most other dyads, suggesting more common signaling strategies.

Importantly, distinct signaling strategies do not imply poorer coordination. For instance, D9 achieved the earliest button presses, and D1 pressed at the smallest distance from the start (see Supplementary Note 6). Therefore, some dyads may have developed individual “dialects” that facilitated more efficient communication.

### 1.4 SHAP-based interpretation of model decisions

The Random Forest model compresses high-dimensional sensorimotor states into target confidence, but like many machine-learning models, its decision process is not directly interpretable. Therefore, we used SHAP (SHapley Additive exPlanations), an explainable AI method that quantifies the contribution of each feature to the model output [39, 40]. Specifically, SHAP decomposes the RF prediction for each sample into a baseline and individual feature’s contribution. The resulting SHAP values show how each feature affects the model’s prediction, with positive values increasing the RF confidence.

The mean absolute SHAP values for each feature and target provide an overall ranking of feature importance (Fig. 5A). Across all targets in Scene 2, table position was the most important feature, followed by table velocity direction and leader’s forces. Table velocity contributed primarily to *T* 3 and *T* 4, which were located in a narrower region. The remaining variables had less influence. Similar results were observed for Scene 1 (see Supplementary Note 4.2).

**Figure 5:**
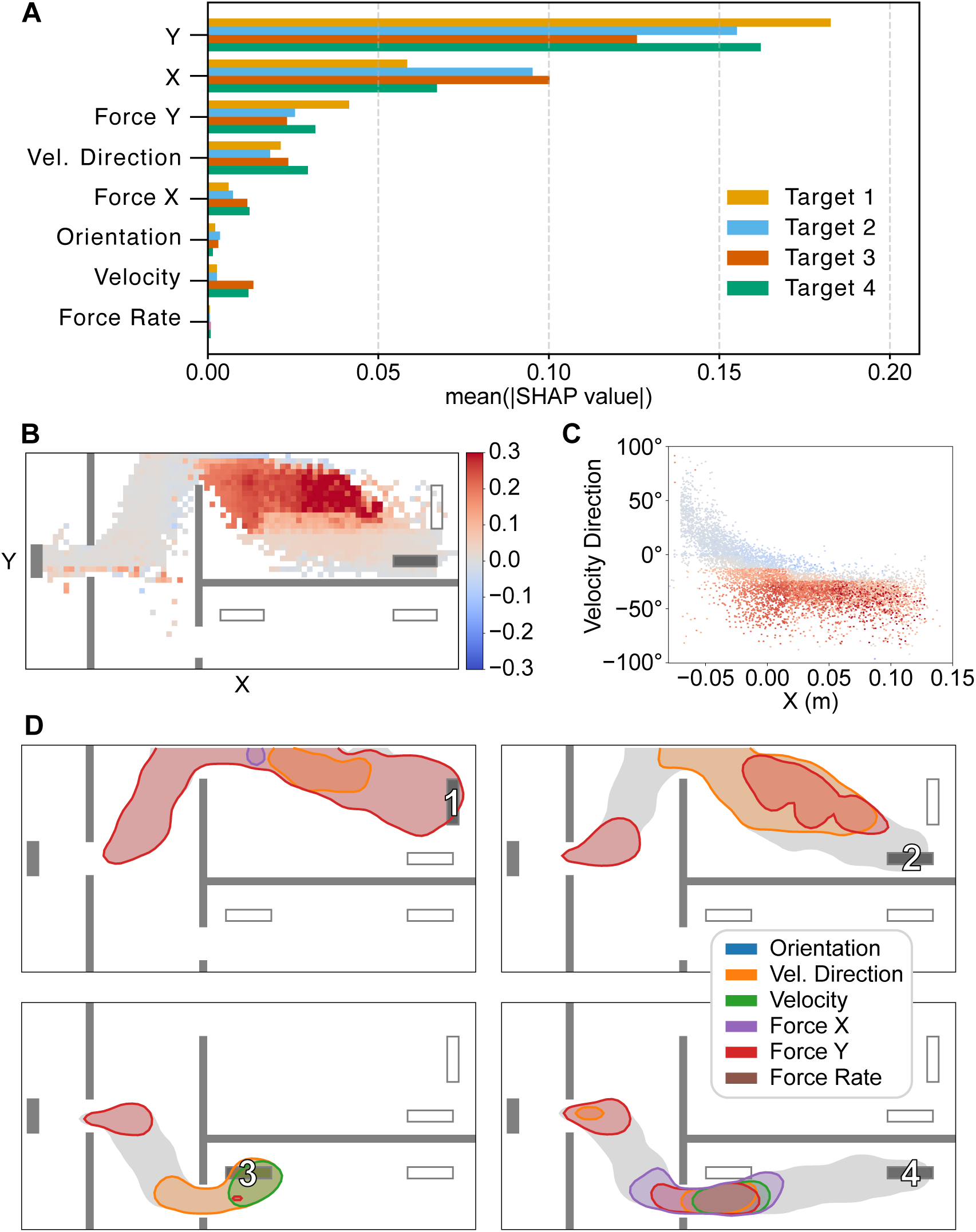
SHAP analysis explaining the RF predictions and feature contributions in Scene 2. **A.** Mean absolute SHAP values for each feature and target, showing the overall importance of each feature. **B.** Spatial distribution of SHAP values for table velocity direction for *T* 2, computed by discretizing the workspace into spatial bins and computing the 95th percentile within each bin. Color indicates the signed SHAP value, with positive values increasing the model’s confidence. **C.** Dependence of the contribution of velocity direction on horizontal position for *T* 2. Points show samples for *T* 2 plotted by table *x* position and velocity direction, with color indicating the signed SHAP value of velocity direction. **D.** Spatial regions where each feature substantially increased the model’s confidence for each target. Contours indicate regions where the 95th percentile of the signed SHAP value exceeded 0.1 (probability units), color-coded by feature. Table position was excluded to highlight where other kinematic and kinetic features contributed. The gray shading indicates the overall trajectory distribution for each target.

The spatial distribution of SHAP values further illustrated how feature contributions varied across the workspace. As an example, Fig. 5B shows the spatial distribution of SHAP values for table velocity direction for *T* 2 in Scene 2. Positive SHAP values were concentrated in the upper part of the top-right room, indicating where this feature was most informative. This agrees with our empirical expectation: before the second passage, velocity direction cannot distinguish between *T* 1 and *T* 2, whereas in the lower part of the top-right room, position is a sufficient cue and velocity direction adds little additional information. Additionally, positive SHAP values were associated with negative angles below *−*20*^◦^* (Fig. 5C), indicating that downward-directed motion in this region increased the model’s confidence in *T* 2. This analysis reveals how each variable should be modulated to make the movement more legible.

Summarizing across features, the overlaid SHAP map shows where each feature contributed across the workspace (Fig. 5D). Leader’s vertical force contributed prominently after the first passage and helped distinguish whether the intended target lay in the upper or lower part of the scene. In the top-right room, where targets were farther from the passage in a more spacious room, the model relied mainly on the leader’s vertical force and the table velocity direction. In contrast, the bottom-right room was more spatially constrained, especially for *T* 3, which was close to the passage. For this target, table velocity and velocity direction contributed more strongly. For *T* 4, the trajectory initially passed near *T* 3, and the model relied on a combination of multiple force and velocity cues. Once the table moved beyond *T* 3, position alone became sufficient to distinguish *T* 4. Overall, these SHAP maps showed that RF relied on different cues depending on the local geometry, reflecting how humans adjusted signaling across sensorimotor variables to the local context. Similar results were obtained for Scene 1 (see Supplementary Note 4.2).

Together with the mechanical work results, these analyses suggest that dyads flexibly combined kinematic and force cues in a context-dependent manner to increase movement legibility, thereby communicating efficiently and effectively without energy-costly detours.

### 1.5 Modeling the follower’s decision-making process

In the *Button Press* condition, the uninformed participant gradually accumulated evidence from the ongoing sensorimotor signals and pressed a button once they identified the target, providing empirical decision times. The broad distribution of table positions at the time of the button presses (Fig. 6A, Supplementary Note 5) reflected a large variability in signaling strategies, cue interpretation, and reaction time. The relatively rare incorrect choices (marked as *×*) tended to occur in spatially ambiguous regions, where samples associated with different targets overlapped.

**Figure 6:**
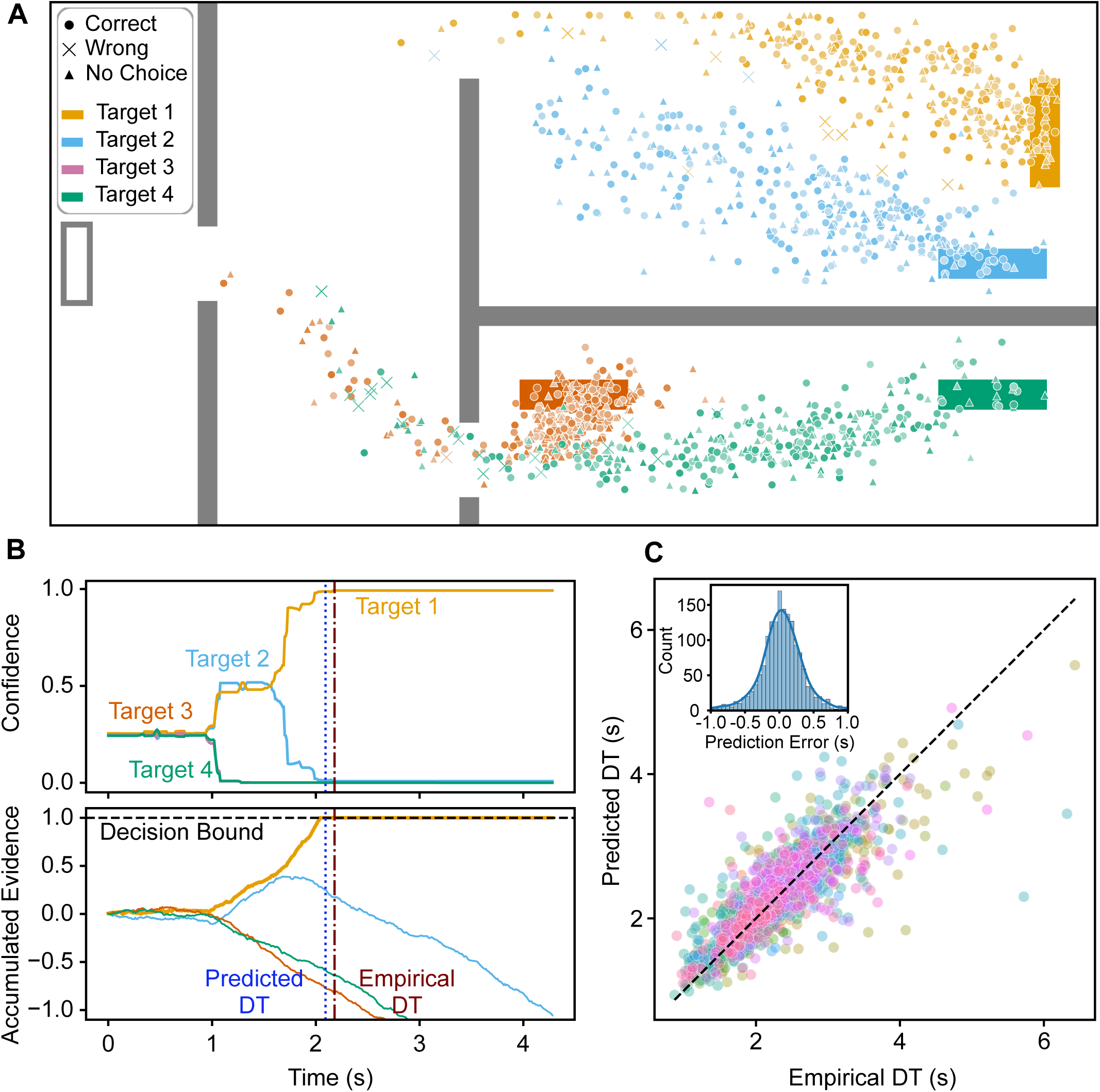
Modeling human decision-making process using a racing DDM. **A.** Table positions at the moment of the button press across all participants for Scene 2. Lighter and darker shaded points indicate earlier and later trials, respectively. **B.** Example trial. Top: Confidence traces predicted by the RF model for each target. Bottom: Corresponding evidence accumulation trajectories in the DDM. Vertical lines mark the empirical (brown) and the predicted decision time (blue) corresponding to when a target’s accumulated evidence first crossed the decision bound (horizontal dashed line). **C.** Decision time prediction. Comparison between the predicted and empirical decision time (DT) for every button press trial, color-coded by participant. Inset: Distribution of prediction errors, overlaid with a kernel density estimate.

To model this decision-making process, we applied the widely used Drift-Diffusion Model (DDM) framework. In its classical form, DDM assumes a one-dimensional evidence signal. Thus, it is not directly applicable to our task, where signals are high-dimensional and must first be interpreted before being treated as evidence. Therefore, we combined the DDM with the RF decoder. Specifically, we used the RF to map the high-dimensional feature state to one-dimensional time-varying confidence traces for each target. These traces were then used as inputs to the DDM, which accumulated positive evidence when confidence for a target exceeded chance level (0.25 for four targets), and negative evidence otherwise. We implemented a racing DDM in which four accumulators, one for each target, accumulated evidence in parallel. For each trial, the first accumulator to reach the decision bound predicted the target choice of the uninformed participant, with its crossing time predicting the decision time. One racing DDM was fit for each participant using the Button Press trials in which that participant was uninformed.

A representative trial illustrates the DDM process, with both the confidence inputs and the corresponding accumulated evidence traces (Fig. 6B). Early in the trial, confidence stayed near chance level for all targets, leading to near-zero evidence accumulation. As informative cues emerged, RF confidence increased for the target (*T* 1) and decreased for the decoys. Correspondingly, *T* 1’s accumulator rose and crossed the positive bound, whereas competing accumulators drifted downward. The predicted decision time closely matched the participant’s button press time.

Across participants, the racing DDM predicted the participant’s chosen target with an accuracy of 94.9%*±*5.1% (mean*±*SD). Predicted decision times closely matched empirical ones, with points clustered around the unity line (Fig. 6C). Prediction errors were small and approximately unbiased (0.02*±*0.37 s; Fig. 6C, inset), indicating little systematic bias. That is, the DDM could accurately predict both the selected target and decision time of the participants for each trial, suggesting that the RF model effectively captured the target-relevant information carried by the sensorimotor signals.

Decision-making models such as the DDM provide a mechanistic account of how choices are formed over time, offering insight into the latent computations that give rise to behavior. By combining the RF model with a racing DDM, we extended evidence accumulation modeling to high-dimensional sensorimotor signals, in which the relevant evidence is not directly available but must first be inferred. The high prediction accuracy further supports the validity of the machine learning decoding of the sensorimotor strategies.

## 2 Discussion

Physical collaboration often requires partners to combine their available information and coordinate their actions despite unequal access to task-relevant information. In this study, we investigated how humans with asymmetric information convey intent through sensorimotor signals during joint object manipulation. We introduced a unified analysis pipeline that bridges high-dimensional behavior with cognitive decision-making: decoding intent from sensorimotor signals, extracting the context-dependent semantics of specific kinematic and kinetic features, and modeling the evidence accumulation process. Through these analyses, we demonstrate that intent is not conveyed by a single channel, but through the flexible modulation of multimodal kinematic and haptic cues. Critically, increasing legibility did not necessarily come at the cost of transport efficiency. Humans selectively modulated the sensorimotor features that were most informative within the local task geometry, without increasing total mechanical work. Moreover, while most dyads exhibit a shared signaling consensus, we observed the emergence of dyad-specific “dialects” that potentially yielded more efficient communication.

Prior works have often studied legibility in joint action through visual cues, such as the trajectory shape [16, 22], leaving the contribution of haptic signals to legibility less explored. In contrast, our results show that intent is conveyed through integrated multimodal cues that combine visual and haptic signals. Importantly, these communicative cues are embedded within actions that must remain pragmatic for task completion. Within this constraint, dyads flexibly shifted which features they relied on, or combined multiple cues, depending on the local workspace geometry and which decoys were currently plausible. This context-dependent multimodal signaling may have allowed dyads to make their actions legible without additional energetic cost.

Several studies have proposed that deviations from efficient movement should be a core mechanism for intent communication [16, 25]. Consistent with this idea, observers often interpret inefficient actions as attempts to convey information [41, 42, 43]. However, this view has largely been developed in tasks with limited action redundancy, where only a small number of efficient alternatives are available to achieve the goal. In such settings, communicative modulation may necessarily appear as deviations from efficiency. By contrast, in our joint object manipulation task, participants increased movement legibility without compromising efficiency. Joint object manipulation involves multimodal, high-dimensional sensorimotor signals, creating natural redundancy in how the task can be completed. Participants could therefore select more legible actions from among similarly efficient alternatives, and introduce communicative modulations that either contributed to task completion or carried little additional energetic cost. This may have allowed dyads to communicate intent without compromising dyadic efficiency.

We found a shared consensus on sensorimotor signaling strategies across the majority of dyads. This ‘universal language’ could arise from the task’s physical constraints as well as participants’ past experiences in joint actions, reflecting shared principles through which communicative meaning is embedded in, and inferred from, movement. Such shared principles may provide a foundation for human physical collaboration, allowing even unfamiliar partners to rapidly establish mutual understanding and coordinate their actions. However, as dyads repetitively worked together, we observed the emergence of dyad-specific ‘dialects’ characterized by systematic deviations from the general strategy. These dialects potentially enabled pairs to improve communication. This suggests that sensorimotor communication is both robust and adaptive, relying on broadly shared baseline strategies that can be further specialized as partners converge on dyad-specific features.

Importantly, we introduced in this paper a framework to decipher the high-dimensional sensorimotor communication by integrating (i) the predictive power of machine learning (Random Forest), capturing statistical regularities in sensorimotor communication, (ii) the transparency of explainable AI (SHAP), quantifying the relative contribution of each feature to the information carried by sensorimotor signals, and (iii) models of human decision making from cognitive neuroscience (Drift-Diffusion Models), simulating the process of evidence accumulation and decision-making. A challenge in computational modeling of human behavior is the tension between model complexity and interpretability: more complex models can capture richer behavioral structure, but are often harder to interpret. Traditional computational approaches in motor control often favor compact, interpretable models, such as Bayesian inference or optimal control frameworks, which offer deep mechanistic insights and clear conceptual narratives. However, these models can become computationally costly, even for seemingly simple and constrained tasks. For example, action understanding has been formalized as Bayesian inverse planning [44], and later extended to Bayesian theory-of-mind models [45], but scaling such models to richer observation spaces and continuous movement dynamics remains challenging. The difficulty is especially pronounced in sensorimotor signaling, where the meaning of an action depends on a complex interplay between multiple coupled variables, the local geometry, and uncertain predictions of the partner’s responses.

To address these limitations, modern work in computational neuroscience increasingly integrates machine learning models to handle complex and large-scale data [46, 47]. Our framework leverages Explainable AI as a scalable solution that can handle high-dimensional behavioral data while opening the “black box” to reveal internal model dynamics. We decoded sensorimotor signals and localized the contributions of specific features across diverse contexts. Moreover, our pipeline extended Drift-Diffusion Models from accumulating evidence over one-dimensional stimuli to high-dimensional sensorimotor states. In this way, we provide a unified computational framework for studying how humans communicate through sensorimotor signals.

Robotics research often treats generating legible motions as an optimization problem to maximize a legibility function [48], where legibility is defined with hand-specified efficiency and rationality rather than learned from human-human collaboration. As a result, optimization can yield motions that are legible but not human-like, deviating from the conventions that humans naturally use when coordinating with each other. We argue that progress toward human-like legibility requires studying sensorimotor communication in human–human experiments, where signaling emerges in its most natural form with physical coupling and multimodal feedback. By grounding legibility in empirically observed patterns, our approach provides a reference for robots to communicate through the shared object using the same channels humans rely on, and it supports a unified framework to infer intent from movement, and to shape movement given intent.

It should be noted that communication is inherently bilateral [49], yet our measurements and analyses primarily characterize how the informed partner’s movement and forces reveal intent, with limited access to how the uninformed partner contributes to the exchange. In practice, followers may signal uncertainty, confirm understanding, or actively probe. Such feedback loops are only partially captured when the analysis is centered on leader-driven cues. Furthermore, while the Random Forest and SHAP analyses identify predictive features, they do not recover a unique “true” signal. Since many kinematic and force variables are mechanically coupled, correlated features can trade off in importance estimates. We alleviated this by selecting relatively independent features, but residual coupling is unavoidable in physical interaction and can vary across workspace regions. Finally, our paradigm focuses on one type of information asymmetry, where one partner has more information than the other. However, real collaboration may involve more dynamic information contexts, for instance with an asymmetry that emerges only mid-trial, or interacting partners having different task-relevant knowledge.

Beyond human-human collaboration, learning sensorimotor communication presents a unique challenge for human-robot interactions. Unlike verbal communication, it is inherently flexible and adaptive and lacks a stable mapping from a particular force–motion pattern to its meaning. This makes it challenging to generate comprehensive descriptions for an AI or a robot to learn from. Recent progress in training robots from large-scale video data [50] is structurally limited for physical collaboration, where haptic signals such as interaction forces and limb stiffness are critical, but not visually observable. While robots could learn by directly interacting with humans, collecting such data is resource-intensive and may pose safety risks during physical interaction. More importantly, humans are extremely adaptive and may rapidly infer a robot’s limitations before it can learn, which could lead them to switch to simplified or exaggerated strategies. The robot would thus end up learning downgraded and suboptimal patterns.

In contrast, by grounding legibility in empirically observed behavioral patterns, our approach provides a usable reference for deciphering human-human interactions. The decoded communication patterns could inform the design of robots that communicate intent in a human-compatible way, with potential applications in physical assistance, rehabilitation, joint training, and sensorimotor augmentation [51].

## 3 Methods

### 3.1 Participants and materials

#### Participants

The experimental protocol was approved by the Ethics Committee of the Technical University of Munich (approval number 763/20 S-KH). Thirty-four right-handed participants (17 dyads; age range 20–35 years; 23 females) were recruited. Written informed consent was obtained from each participant prior to the experiment. All participants were neurologically healthy, reported no motor impairments of their right arm, and were naive to the purpose of the study. Handedness was assessed using the Edinburgh Handedness Inventory [52].

#### Experimental setup

Pairs of participants were seated side by side at a table, separated by an opaque curtain (Fig. 2A). Each participant viewed the virtual environment on a screen in front of them and operated a 3-DoF haptic device (Phantom Premium 1.5 HF, 3D Systems) with their right hand. The haptic device provided force feedback, with the peak force limited to 8 N for safety. End-effector motion was virtually constrained to a vertical plane parallel to the screen (stiffness 200 N*/*m). Visual rendering ran at 60 Hz, and haptic feedback was presented at 1000 Hz. End-effector position and force were stored at 1000 Hz for offline processing.

#### Virtual environment

The virtual apartment environment was simulated in 3D using Chai3D and Open Dynamics Engine [53, 54]. We rendered a workspace (40 cm *×* 20 cm) containing a table (4 cm *×* 1 cm, 0.5 kg), four possible targets with the same geometry as the table, and walls that partitioned the workspace into four rooms. Adjacent rooms were either connected by a 3 cm-wide passage or separated by walls. We designed two scenes with the same wall layout but different target layouts (Fig. 7A): *T* 1, *T* 2, and *T* 4 had the same locations in both scenes, whereas *T* 3 differed. As a result, Scene 1 had three potential target locations at the top and one at the bottom, while Scene 2 had two potential target locations on each side. Consequently, the workspace contained rooms with varying ambiguity (i.e., one, two, or three candidate targets) and varying spatial constraints and space for signaling. Target visibility depended on the experimental condition (see *Methods*–*Experimental conditions*).

**Figure 7:**
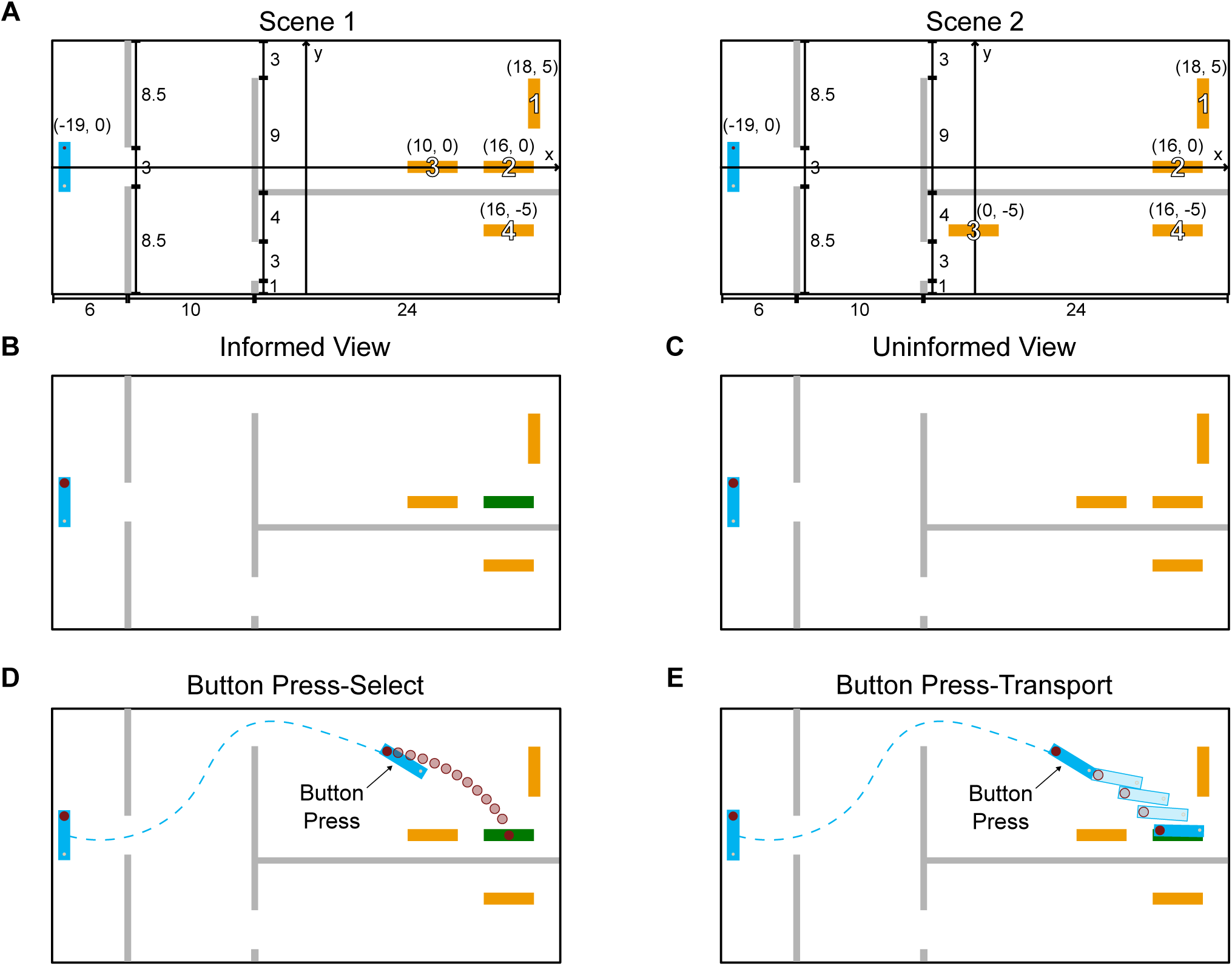
Experimental Protocol and Setup. (A) Scene layouts of Scene 1 and Scene 2 showing the starting position (blue) and the four potential target locations. Dimensions and coordinates are expressed in centimeters. (B) The screen as seen by an Informed participant, where the target is green and decoys are orange. (C) The screen as seen by an Uninformed participant, where all potential targets appear identical in orange. (D–E) Illustrations of the two trial types in the Button Press condition. The task requirements differ in these two conditions after the button press: (D) Button Press–Select, the uninformed partner selects a target with a cursor; (E) Button Press–Transport, the dyad physically moves the table to the target.

The two partners in an experiment jointly controlled the table object at its two ends (Fig. 2A). Each hand position was connected by a virtual spring (stiffness 800 N*/*m) to a control point located on the table centerline (parallel to the long axis), positioned 0.5 cm from the edge. The haptic devices rendered forces arising from interactions with the environment and the partner via the table. Both partners viewed a top-down rendering of the scene, and their on-screen motion matched the direction of their physical hand motion. The friction coefficient between the table and the floor was 1.0, and both the table and the walls had a contact stiffness of 300 N*/*m. Upon collision, the wall turned red, a beep was played, and participants felt the resulting contact force through their haptic device.

### 3.2 Experimental paradigm

#### Task and conditions

The experimental task required dyads to collaboratively transport the table object to a target while avoiding obstacles. At the onset of each trial, the table was fixed at the start location and participants moved their cursors to their assigned control points to “pick up” the table. When both cursors were within 2 cm of their assigned control points, each cursor was coupled to the corresponding control point via a virtual spring, so participants could move the table. At the same time, four potential targets appeared, and participants slid the table to the target, while avoiding collisions with the walls or entering decoys. Verbal communication was not allowed. Due to simulated friction, the table could be transported efficiently only when both participants contributed force in coordination, providing a natural incentive to collaborate.

We designed three block conditions: Symmetric, Asymmetric, and Button Press (Fig. 2B). Conditions differed in (i) which participant could visually identify the target and (ii) how the trial ended. In the Symmetric condition, both participants saw the target highlighted in green, while the three decoys were shown in orange (Fig. 7B). In the Asymmetric and Button Press conditions, only one participant was informed of the green target, and the uninformed participant saw four identical orange targets (Fig. 7C). Additionally, in the Button Press condition, the uninformed participant pressed a button in their left hand as soon as they could identify the target. After the button press, either (i) the table was frozen and the uninformed participant selected the target they believed to be correct with their cursor (Button Press–Select condition, Fig. 7D), or (ii) the dyad continued transporting the table to the target (Button Press–Transport condition, Fig. 7E). These two trial types were randomized, and the dyad only learned the current trial type after the button press.

In the Symmetric, Asymmetric, and Button Press–Transport trials, trials ended when the table reached the target, defined as the center of the table being within 2 cm of the target position and its orientation within 20*^◦^* of the target orientation. In the Button Press–Select trials, trials ended with the uninformed participant’s target selection.

#### Scoring

At the end of each trial, both partners were shown the team score. In Symmetric, Asymmetric, and Button Press–Transport trials, this score was based on completion time from pickup to successful placement. The dyad received 5 points for completion time under 4 s; 1 point was deducted for each additional second, to a minimum of 0 points. In Button Press–Select trials, if the correct target was selected, the score was based on the time from pickup to button press: the dyad received 8 points for button press time within 1 s, with 1 point deducted for each additional second to a minimum of 0 points. An incorrect target selection yielded 0 points. Participants were instructed to avoid collisions with walls and entry into decoys, but these did not affect the score.

#### Experimental Protocol

Each dyad completed a total of 13 blocks, with 3 Symmetric, 6 Asymmetric, and 4 Button Press blocks. The experiment followed a fixed block sequence shown in Fig. 2C to control for learning and role-specific effects. In the Asymmetric and Button Press blocks, the right-seated participant was visually informed of the target and the left-seated participant was uninformed in blocks 2–6; roles were reversed in blocks 8–12.

Each block contained 40 trials, with each target in each scene presented five times. To evenly distribute targets within a block, 40 trials were arranged into five sets of eight trials, with each target in each scene presented once per set in randomized order. In Symmetric and Asymmetric blocks, all trials within a block followed the same information structure and trial protocol. In Button Press blocks, each target appeared in two Transport and three Select trials (16 Transport and 24 Select trials per block); Transport and Select trials were randomly distributed and only revealed after the button press. This design ensured two requirements simultaneously: the informed participant’s movements needed to be informative early and physically feasible for transport. Participants were informed of the block condition at the start of each block, but not of the trial order. Before the main experiment, participants completed familiarization trials. Each participant completed five trials in each condition (Symmetric, Asymmetric, Button Press) in each role (informed and uninformed, where applicable) using unique target layouts. Button Press familiarization included both trial types.

### 3.3 Data analysis

Kinematic and dynamic data were low-pass filtered (Butterworth, 20 Hz cut-off frequency, tenth order) before computing the reported metrics. Data analysis was performed with Python 3.8. The *mechanical work* expended by each participant was computed as

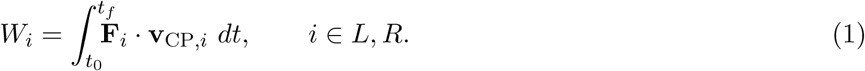

where *i* denotes the left- (L) or right-seated (R) participant, **F***_i_* is the 2D force applied by participant *i* at their control point on the table, and **v**_CP_*_,i_* is the 2D velocity of that control point. Total work was computed as *W*_tot_ = *W*_L_ + *W*_R_. The SPARC smoothness reported in the Supplementary materials was computed using the codes provided in [55].

### 3.4 Statistical analysis

Several statistical approaches were implemented using the pingouin and mne.stats package (Python 3.8). All pairwise tests were two-tailed, with a significance level of *α* = 0.05.

#### Block-level metrics

Completion time, expended mechanical work, and work share were compared across experimental conditions using data from the final blocks of each condition (Blocks 5–7 and 11–13). Normality was assessed with Shapiro–Wilk tests [56]. Main effects of condition were tested using repeated-measures ANOVA. If sphericity was violated, a Greenhouse–Geisser correction was applied. Following a significant main effect, post-hoc pairwise comparisons were performed using paired *t*-tests with Holm–Bonferroni correction. For significant post-hoc tests, we report Cohen’s *d* as a measure of the effect size. Statistical significance is indicated as follows: *(*p <* 0.05), **(*p <* 0.01), and ***(*p <* 0.001).

#### Signaling trajectories

To determine where trajectories of signaling variables (e.g., path, orientation, velocity, force) diverged across conditions, we used a non-parametric cluster-based permutation test for within-subject designs [57]. Table position *x* was discretized into 0.01 m bins, and each signal was averaged within bins for each participant, separately for each scene and target. At each *x* bin, a repeated-measures *F* -statistic comparing the three conditions was computed. Contiguous bins exceeding the cluster-forming threshold (*p <* 0.05) were grouped into clusters, and cluster significance was assessed by comparing the summed *F* -values against a null distribution generated from 5000 random permutations of condition labels. Clusters were considered significant if the cluster-level *p*-value was *<* 0.05.

#### Confidence trajectories

To compare confidence increase across conditions, table position *x* was normalized to a unit interval between the second passage (*x* = *−*0.04) and the center of the intended target, then discretized into 2.5% spatial increments. Pairwise non-parametric cluster-based permutation tests [57] were then conducted between conditions, with Holm–Bonferroni adjustment.

### 3.5 Decoding Sensorimotor Communication

As shown in the Results, sensorimotor communication during joint object manipulation is expressed through multimodal, high-dimensional signals. The interpretation of cues is conditional on both task context and the set of co-occurring cues. Because cue effects can interact nonlinearly, their semantics are not additive. We therefore adopted an explainable machine learning approach to relate the sensorimotor signals to the inferred target. This approach comprised two steps: (i) Random Forest models integrated the multidimensional features and produced a probability distribution over candidate targets, and (ii) SHapley Additive exPlanations (SHAP) quantified the contribution of each feature to the model output, enabling interpretation of which signals supported each target prediction and how signals can be shaped to optimize legibility.

#### 3.5.1 Decoding intended target from multimodal cues using Random Forest

In this task, the informed participant conveys the intended target through sensorimotor cues, so the cue interpretation can be formulated as a classification problem. We used a Random Forest classifier [58] because ensembles of decision trees can model nonlinear decision boundaries and feature interactions, are well-suited to high-dimensional sensorimotor signals, and often perform effectively with small-to-moderate training sets.

We trained Random Forest (RF) classifiers using the Scikit-learn library [59] to decode the target from the instantaneous sensorimotor state at each timestep. Input features comprised table position (*x, y*), orientation (*θ*), speed (‖**v**‖), velocity direction (*θ***_v_**), and the leader’s force (*F_x_, F_y_*) and force rate (‖**F͘**‖). The leader was defined as the participant with higher work share in the Symmetric blocks, and as the informed participant in the Asymmetric and Button Press blocks. RF models treated each timestep as a sample, output class probabilities over the four targets, and predicted the target with the highest probability. Data were downsampled to 100 Hz. Train–test splits were performed at the trial level to avoid temporal leakage. Within each block, each target in each scene appeared five times and one trial was randomly held out for testing (20%), and the remaining four trials were used for training.

We trained a general model using data aggregated across all dyads and dyad-specific models using data from each dyad. Models were trained separately for each scene. Hyperparameters were selected by grid search over maximum tree depth 6, 7, 8, 10, 12, 15, 20, 30 and number of trees 50, 100, 200, 300, using 3-fold cross-validation and accuracy as the scoring metric. To balance predictive accuracy and model complexity, we set the maximum tree depth to 10 for the general model and to 8 for dyad-specific models, both using 50 trees.

To visualize how model confidence varied across the workspace, we discretized it into 0.5 cm *×* 0.5 cm bins.

For each bin, we averaged the model’s predicted probability for the target across all samples within that bin.

To compare model similarities, and whether different models produced similar predictions for the same sensorimotor signals, we evaluated each dyad-specific and the general model on a pooled evaluation dataset containing all dyads and computed pairwise difference between models using the Jensen–Shannon (JS) distance between their predicted probability vectors. JS distance was computed at each timestep for each pair of models and then averaged over timesteps and trials. To visualize the JS distances between dyads, a 3D embedding was generated using metric multidimensional scaling (MDS) [60] based on the dissimilarity matrix using the Scikit-learn library [59]. This projection maps the high-dimensional dissimilarity matrix into a 3D Euclidean space, preserving the original distances as closely as possible.

### 3.6 Interpreting model predictions with SHAP

RF models map high-dimensional sensorimotor cues to a confidence value for each target. However, the mapping is not directly interpretable, which prevents assessing how individual signals and their interactions contribute to the model’s prediction. We therefore applied SHapley Additive exPlanations (SHAP) to quantify feature contributions to the RF outputs [39, 40]. SHAP is based on Shapley values from cooperative game theory [61] and quantifies each feature’s average marginal contribution to the prediction. For each sample, SHAP decomposes the model output into an expected value (the baseline) plus additive contributions from individual features. A positive (negative) SHAP value indicates that the corresponding feature increases (decreases) the predicted probability relative to the baseline, and the magnitude reflects the strength of this influence. SHAP values are expressed here in the unit of predicted probability.

We computed SHAP values for the general RF models at each timestep using all participants’ time series data, separately for each scene. Global feature importance was defined as the mean absolute SHAP value per feature, computed for each target and scene. To characterize how feature contributions varied across the workspace, we discretized it into 0.5 cm *×* 0.5 cm bins, spatially aggregated SHAP values within each bin, and summarized their upper range using the 95th percentile. This shows how much the different features could potentially contribute to the model’s confidence across the workspace.

To compare where each feature contributed most strongly, we overlaid “high-contribution” regions for each feature in a single plot. Specifically, for each feature we thresholded the spatially aggregated SHAP map (SHAP *>* 0.1) to obtain a binary mask and then smoothed it with a Gaussian filter (STD=1.5 bins). The contours for each feature were then overlaid in one plot for each target and scene and the features were color-coded.

### 3.7 Drift-Diffusion Models

To model how the uninformed participant accumulated evidence about the target from the evolving sensorimotor cues, we integrated the Random Forest model with a Drift-Diffusion Model (DDM) [62, 63, 64].

The DDM describes a decision process in which noisy evidence is integrated over time until it reaches a decision bound. Classical DDM formulations assume a one-dimensional evidence stream, whereas our sensorimotor signals are high-dimensional and do not directly represent evidence. We therefore used the RF model to map the signals to target-specific probability estimates, and treated these probabilities as momentary evidence. For each target *i ∈ {*1, 2, 3, 4*}*, we defined the momentary evidence as the deviation of the RF output from chance level: *e_i_*(*t*) = *P_i_*(*t*) *−* 0.25. Here *P_i_*(*t*) is the probability predicted by RF at time *t* for target *i*, and 0.25 is the chance level for four alternatives. When *e_i_*(*t*) *>* 0, the model accumulated positive evidence in favor of target *i*, and vice versa.

Because the task involved four alternatives, we implemented a racing DDM with four accumulators running in parallel, one per target. A decision was made when the first accumulator reached the bound *a*, and the corresponding target was taken as the model’s predicted choice:

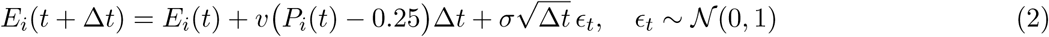

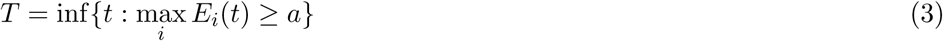

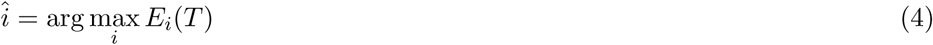

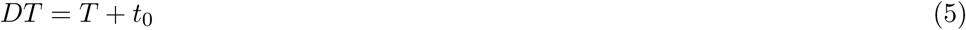

Here, *E_i_*(*t*) is the accumulated evidence for target *i* up to time *t*, *T* the bound-crossing time, ^^^*i* the predicted target and *DT* the predicted decision time including the non-decision time *t*_0_ (accounting for sensory and motor delays). The DDM parameters were the bound *a*, drift gain *v*, diffusion noise *σ*, and non-decision time *t*_0_. We fixed *a* = 1 and fit *v*, *σ* and *t*_0_ using SciPy’s differential evolution optimizer. If the model predicted the correct target, the cost was the absolute error between the predicted and empirical decision time. If the model predicted an incorrect target, we added a fixed penalty of 10 (substantially larger than typical decision-time errors) to strongly penalize incorrect choices.

## Acknowledgments

This work was supported in part by the Deutsche Forschungsgemeinschaft (DFG, project 467042759) and by the European Commission (FETOPEN H2020 899626 NIMA). The project was part of the Imperial-TUM Joint Academy of Doctoral Studies (JADS), where YL was supported by the TUM International Graduate School of Science and Engineering (IGSSE).

## Author contributions

All authors contributed to the conceptualization and methodology of the study. Y.L. developed the software, conducted the experiments and formal analyses, created the visualizations, and wrote the original draft. All authors contributed to the interpretation of the results and provided substantial input on the analyses and visualizations. D.V., E.B., and D.F. reviewed and edited the manuscript. D.F. and E.B. supervised the research and acquired funding.

## Data availability

The data supporting the findings of this study are available from the corresponding author upon reasonable request.

## Code availability

The custom code used to generate the results reported in this study is available from the corresponding author upon reasonable request.

