## Supplementary materials for "Explainable Decoding of Sensorimotor Communication in Joint Object Manipulation"

#### Supplementary Note 1 Learning effects

To examine changes in task performance over the course of the experiment, we analyzed trial-by-trial changes in completion time, movement smoothness and mechanical work across conditions (Fig. S1). The Button Press-Select condition was excluded from this comparison as trials in this condition did not end at the target.

Across trials, participants showed clear improvement in task performance in the early blocks of the experiment, with decreasing completion time and increasing movement smoothness. These improvements gradually plateaued as participants became familiar with the task. After the role switch at Block 8, a similar but smaller adaptation pattern was observed, suggesting that participants adjusted to the new role assignment before performance stabilized again. Combined mechanical work of the dyad remained relatively stable throughout the experiment.

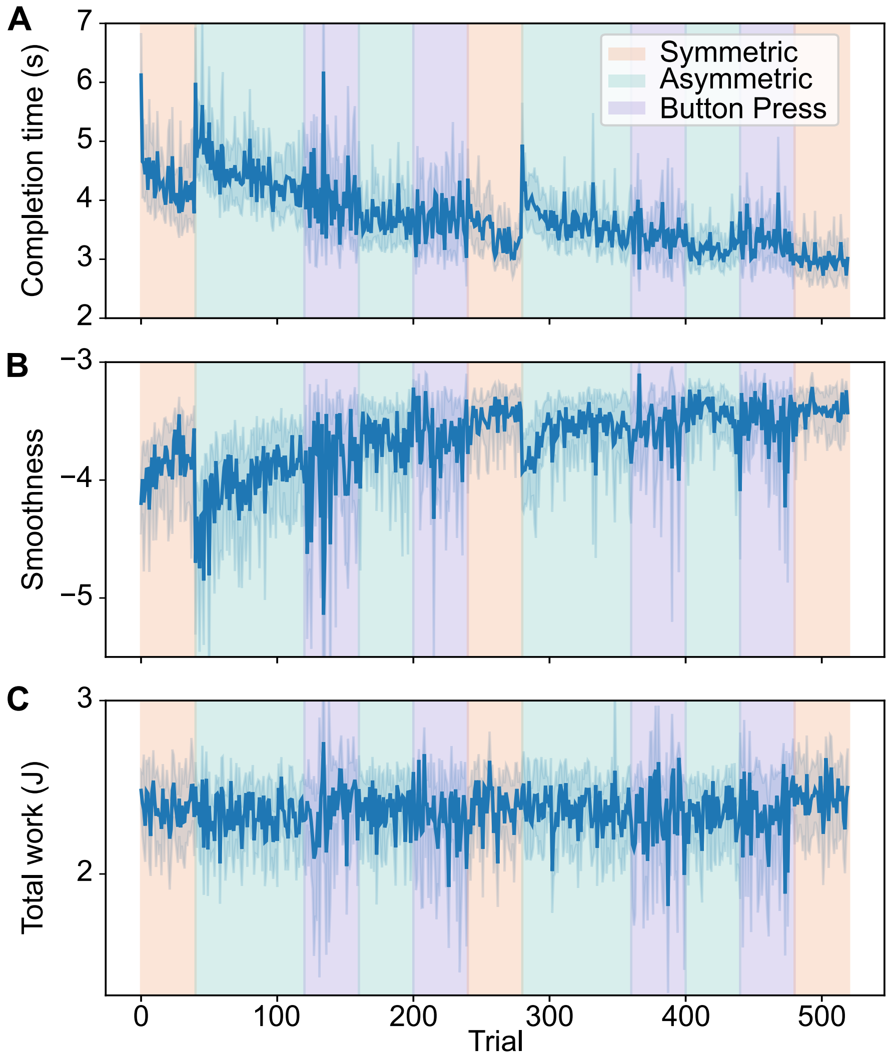


**Figure S1: Learning effects across trials. A.** Completion time. **B.** Movement smoothness quantified using SPARC [1, 2], where larger values indicate smoother movements. **C.** Combined mechanical work of the dyad. Background colors indicate the condition of each block: *Symmetric*, *Asymmetric*, and *Button Press*.

### Supplementary Note 2 Effort-related measures

In Fig. 3B, we showed that participants did not expend additional mechanical work to complete the task in the *Asymmetric* and *Button Press* conditions when compared to the *Symmetric* baseline. To test the robustness of this observation, we computed the total mean force, i.e. sum of mean force magnitude of each partner within a dyad, and the table path length as complementary effort-related measures (Fig. S2).

Consistent with the total mechanical work reported in the main text, total mean force and table path length did not increase in the *Asymmetric* or *Button Press* conditions relative to the *Symmetric* baseline, and even decreased in some comparisons. These results further support that signaling between humans does not necessarily rely on increased effort or larger movement excursions. The corresponding statistical analyses are described in Section 3.4 of the main text.

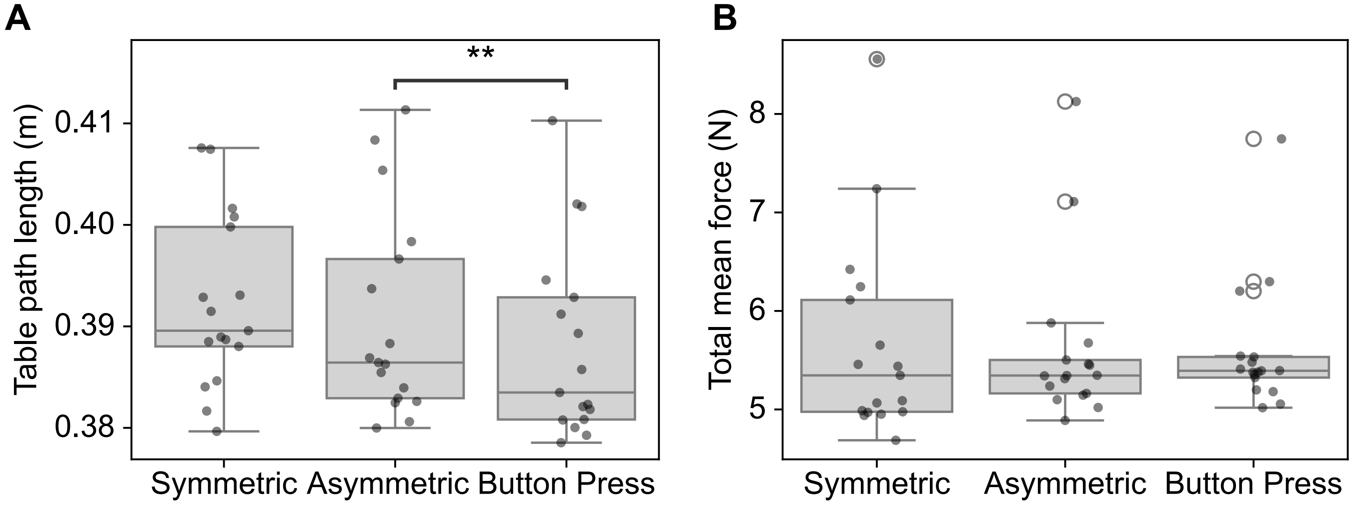


**Figure S2: Complementary effort quantification across conditions.** Metrics were computed using the final blocks of each condition, i.e. blocks 5–7 and 11–13, to mitigate the learning effects. **A.** Table path length, defined as the accumulated translational movement of the table within a trial. **B.** Total mean force, computed by averaging the force magnitude for each participant within a trial and then summing across the two participants in a dyad.

### Supplementary Note 3 Multimodal signaling

In Fig. 3D–G, we showed examples of how sensorimotor signals deviated in the *Asymmetric* and *Button Press* conditions when compared with the *Symmetric* baseline. These deviations may reflect intentional shaping of the sensorimotor signals to make the movement more legible.

Here, we provide a more complete overview of how signaling was embedded across 12 variables (Figures S3–S14). Each figure shows the mean profile of one signal as a function of table position along the $x$-axis for each scene. In all figures, profiles are color-coded by target and shown separately for targets in the upper and lower rooms. Solid, dashed, and dotted lines denote the *Symmetric*, *Asymmetric*, and *Button Press* conditions, respectively. Horizontal bars indicate $x$-ranges with significant differences between conditions according to non-parametric cluster-based permutation tests, color-coded by target. Vertical dashed lines indicate the positions of the constrained passages.

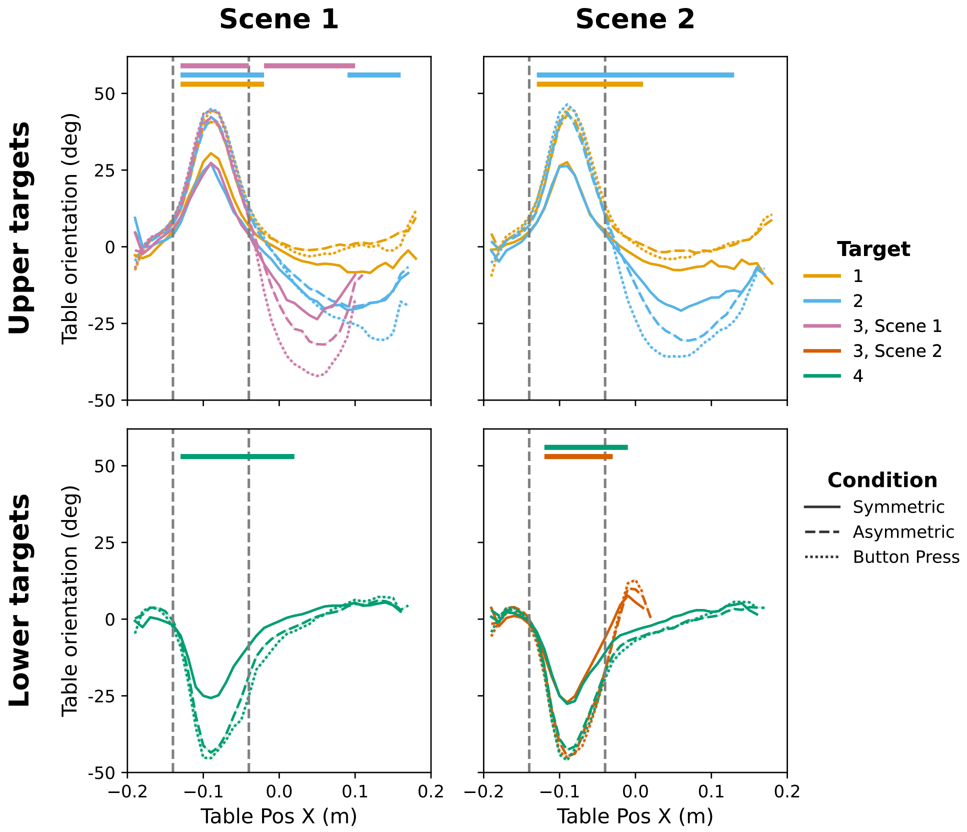


**Figure S3:** Comparison of table orientation profiles across experimental conditions. Positive values indicate counter-clockwise angles relative to the positive $x$-axis.

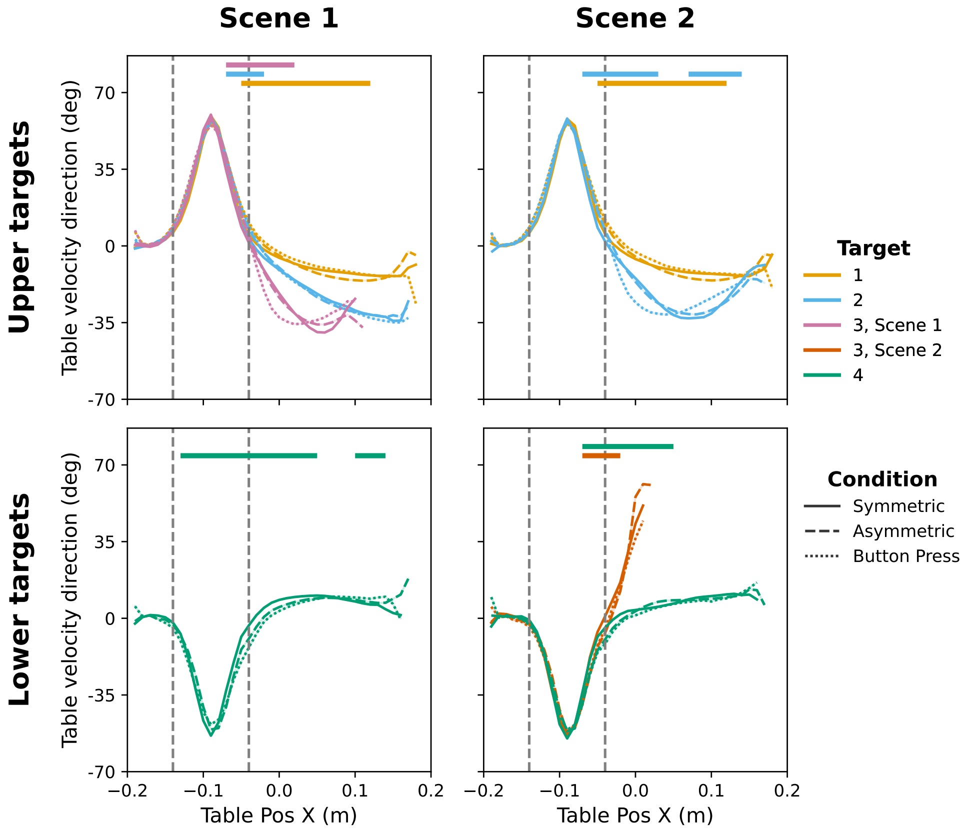


**Figure S4:** Comparison of table velocity direction profiles across experimental conditions. Positive values indicate counter-clockwise angles relative to the positive $x$-axis.

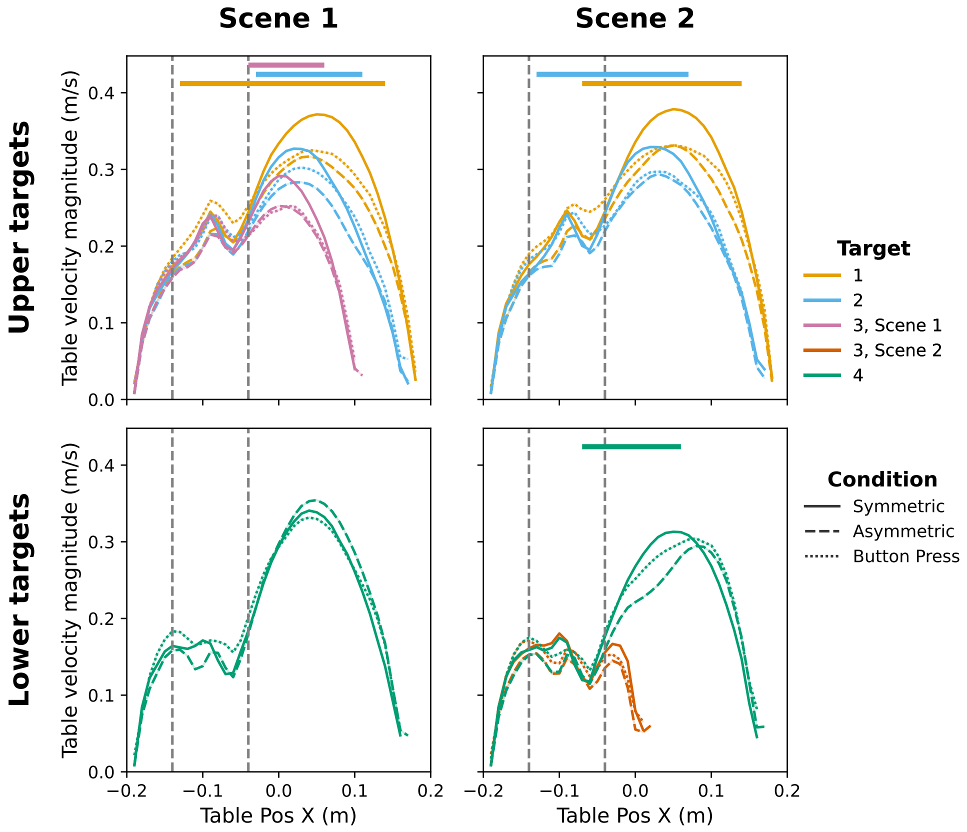


**Figure S5:** Comparison of table velocity magnitude profiles across experimental conditions.

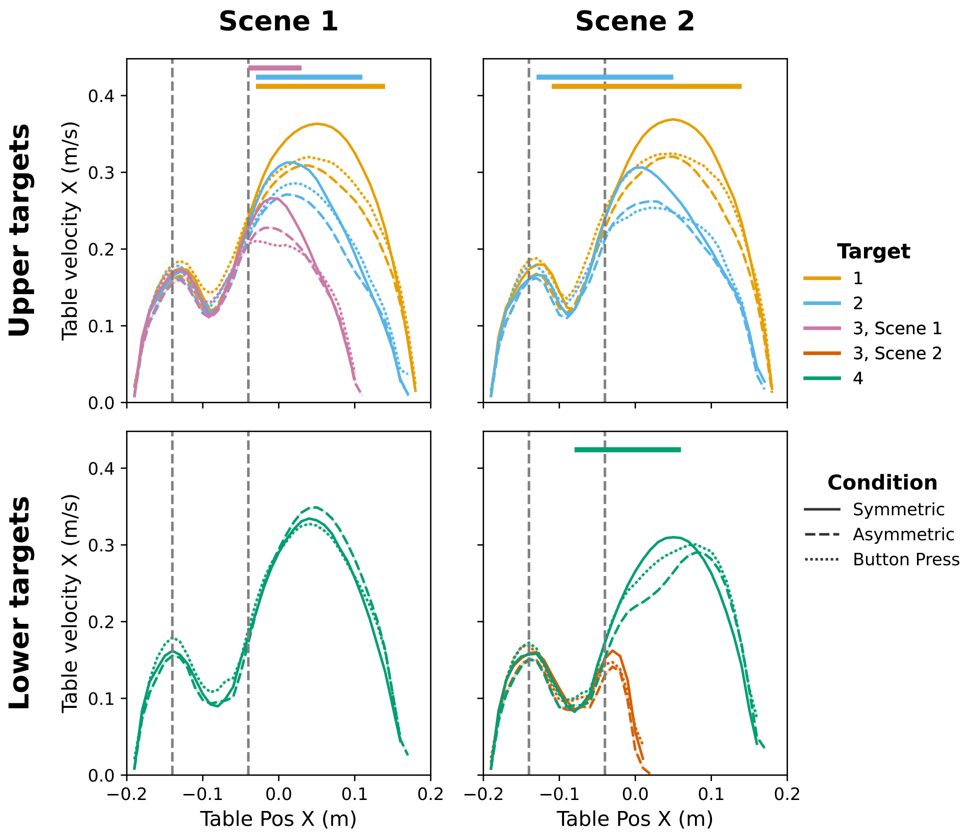


**Figure S6:** Comparison of table velocity $x$-component profiles across experimental conditions.

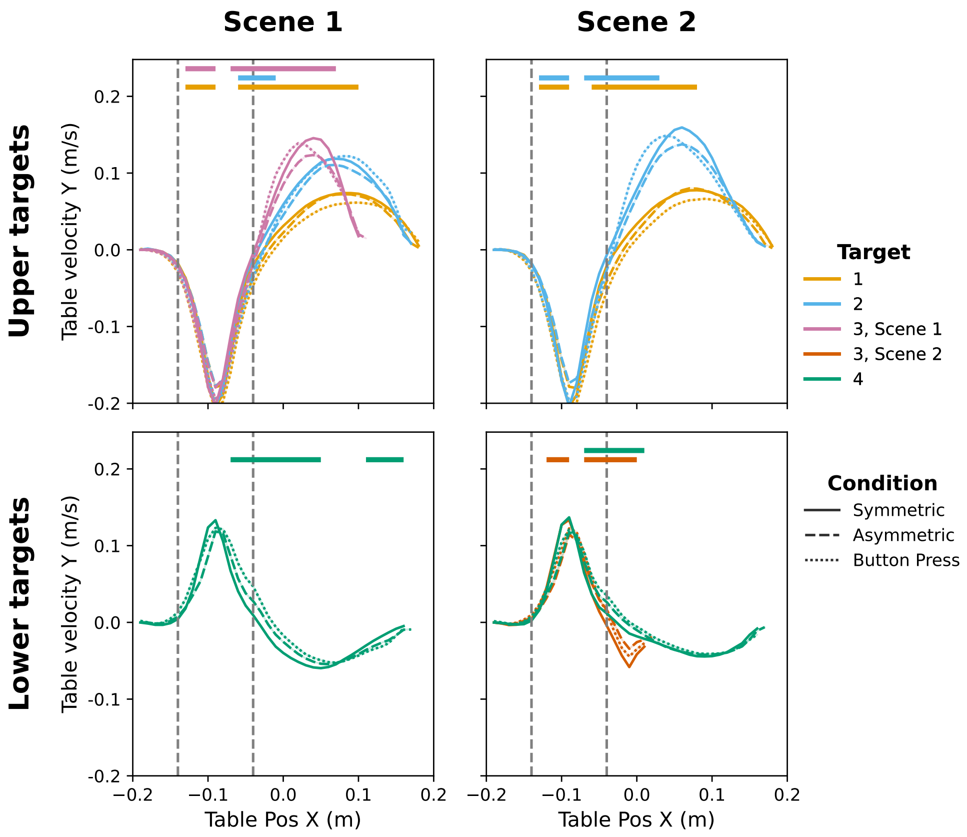


**Figure S7:** Comparison of table velocity $y$-component profiles across experimental conditions.

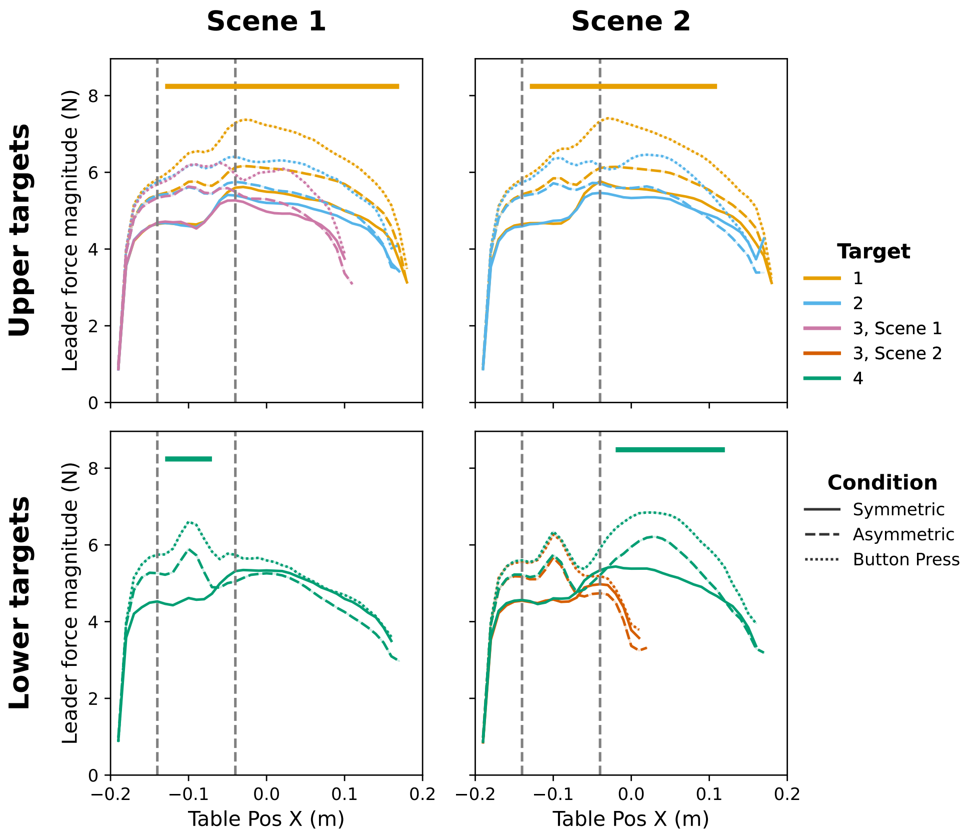


**Figure S8:** Comparison of the leader's force magnitude profiles across experimental conditions.

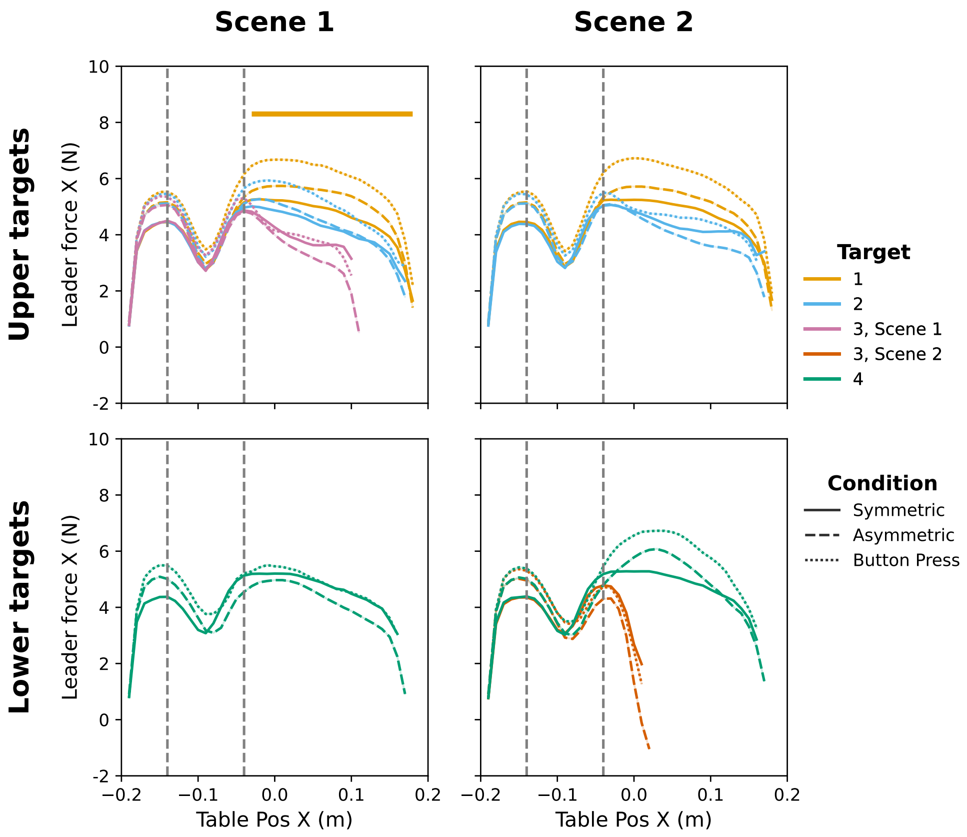


**Figure S9:** Comparison of the leader's force $x$-component profiles across experimental conditions.

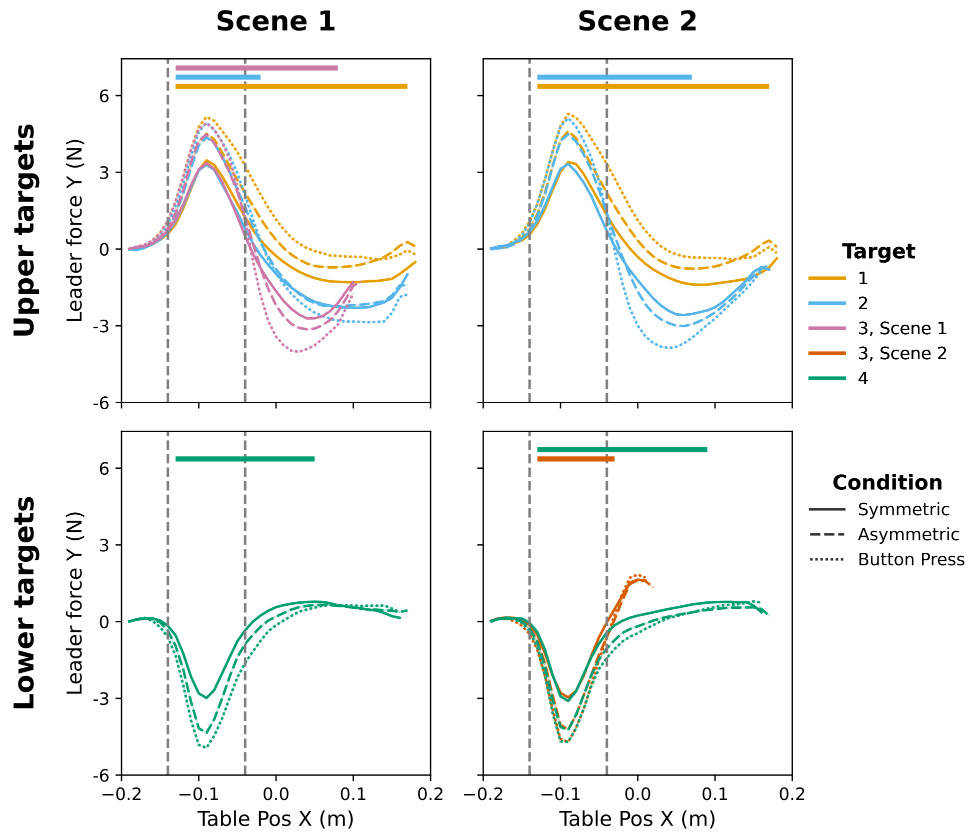


**Figure S10:** Comparison of the leader's force $y$-component profiles along the across experimental conditions.

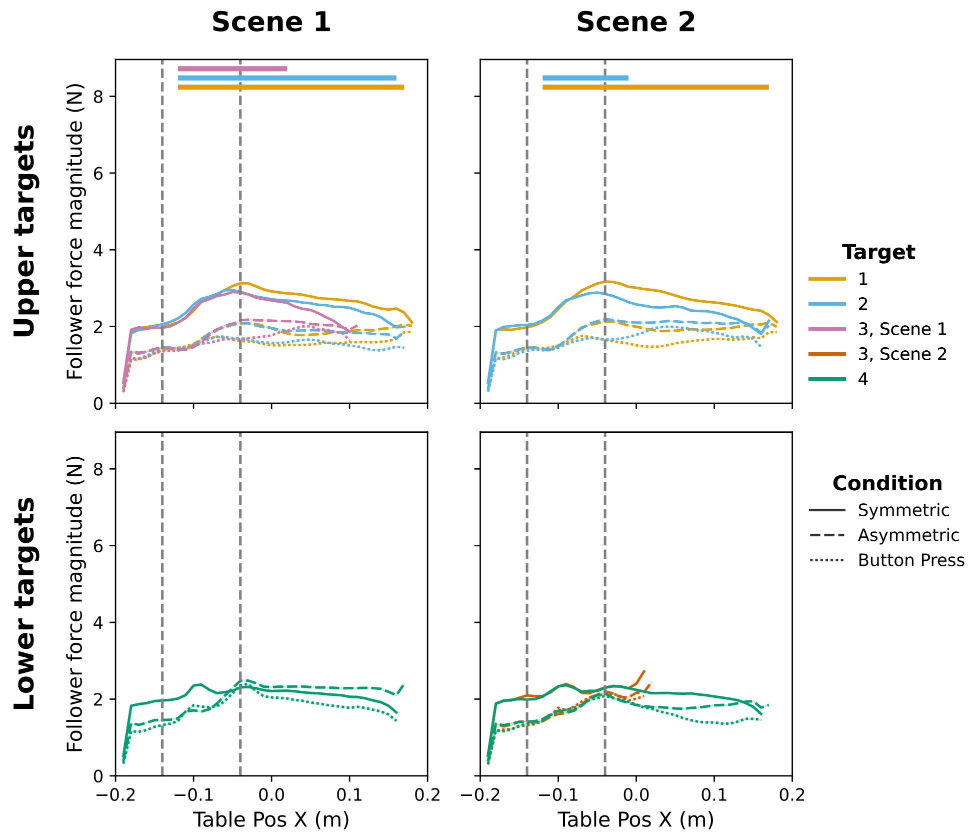


**Figure S11:** Comparison of the follower's force magnitude profiles across experimental conditions.

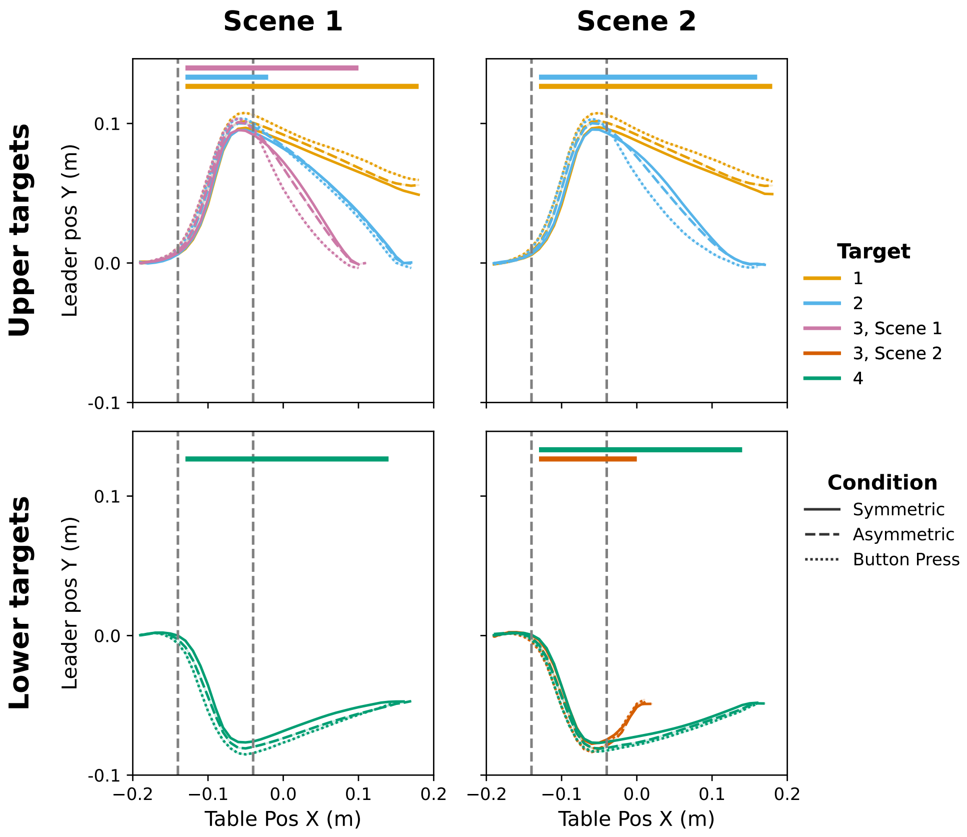


**Figure S12:** Comparison of the leader's $y$-position profiles across experimental conditions.

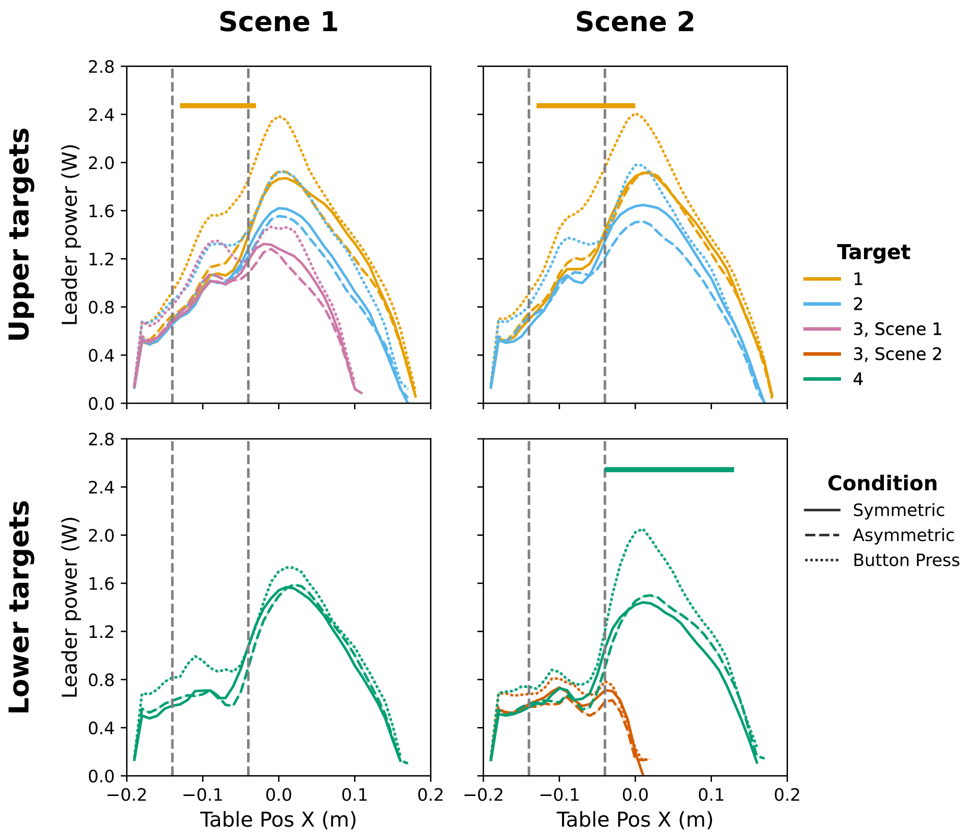


**Figure S13:** Comparison of the leader's power profiles across experimental conditions.

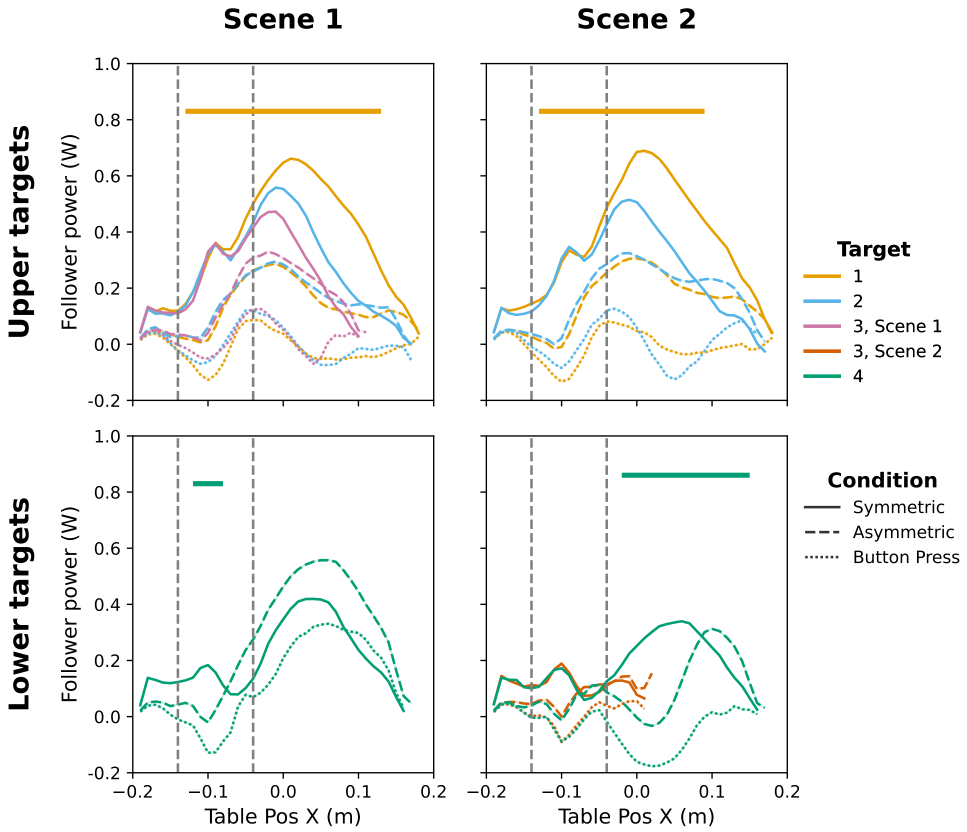


**Figure S14:** Comparison of the follower's power profiles across experimental conditions.

# Supplementary Note 4   Explainable decoding in Scene 1

## 4.1   Random Forest prediction in Scene 1

We showed the spatial distribution of the RF model's confidence in Scene 2 in Fig. 4A. Here, we provide the corresponding confidence distribution in Scene 1. Consistent with Scene 2, confidence was generally higher in regions where the spatial information clearly indicated one target over the others, and lower in areas where trajectories toward different targets overlapped, suggesting that the model's uncertainty reflected the ambiguity of the local task geometry.

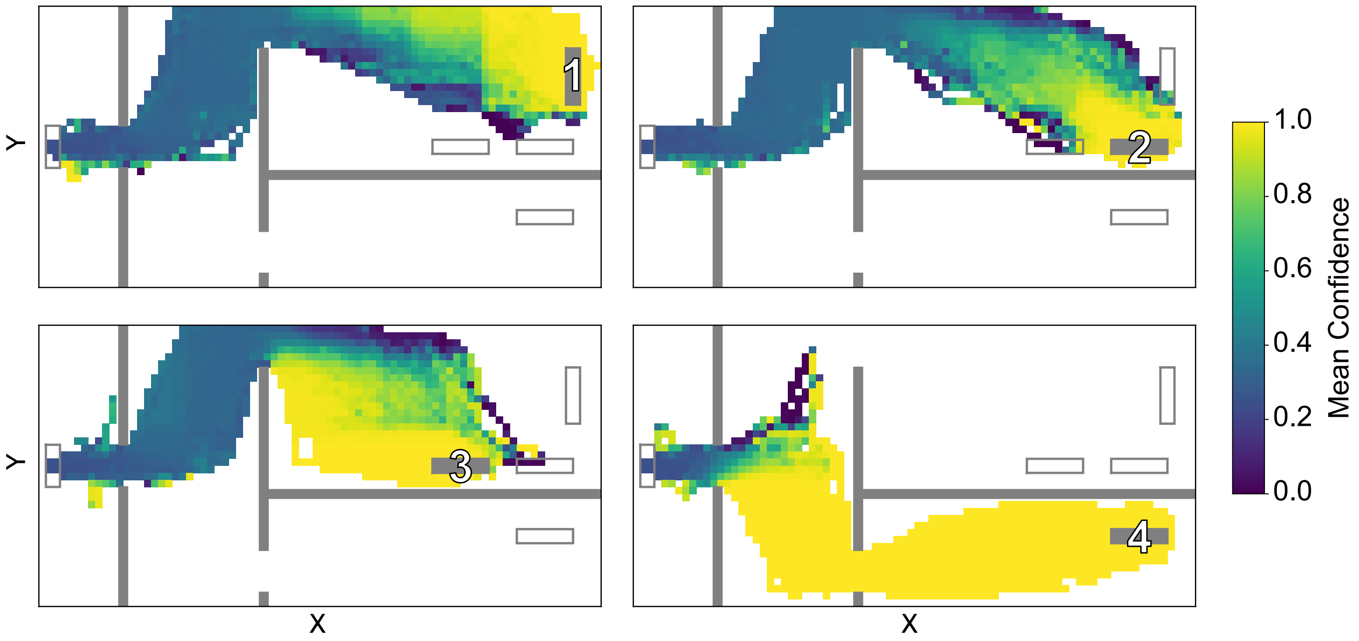


**Figure S15: Spatial distribution of the Random Forest model's confidence in Scene 1**. Mean confidence of the RF model across the workspace is shown separately for each target $T1 - T4$. The workspace was discretized into $0.5 \times 0.5$ cm bins, and color indicates the mean predicted probability assigned to the target within each bin. Higher values indicate regions in which the model predicted the target with greater confidence.

#### 4.2 SHAP analysis in Scene 1

In Fig. 5, we presented the SHAP analysis for Scene 2 to examine how different kinematic and kinetic features contributed to the RF model's predictions across the workspace. Here, we provide the complementary analysis for Scene 1. Similar to Scene 2, the mean absolute SHAP values showed that table position was the dominant contributor across targets. Other features, including the leader's force, table velocity direction, and table orientation, also contributed to the RF's predictions. These results, together with those from Scene 2, show that the RF model relied on context-dependent combinations of sensorimotor features.

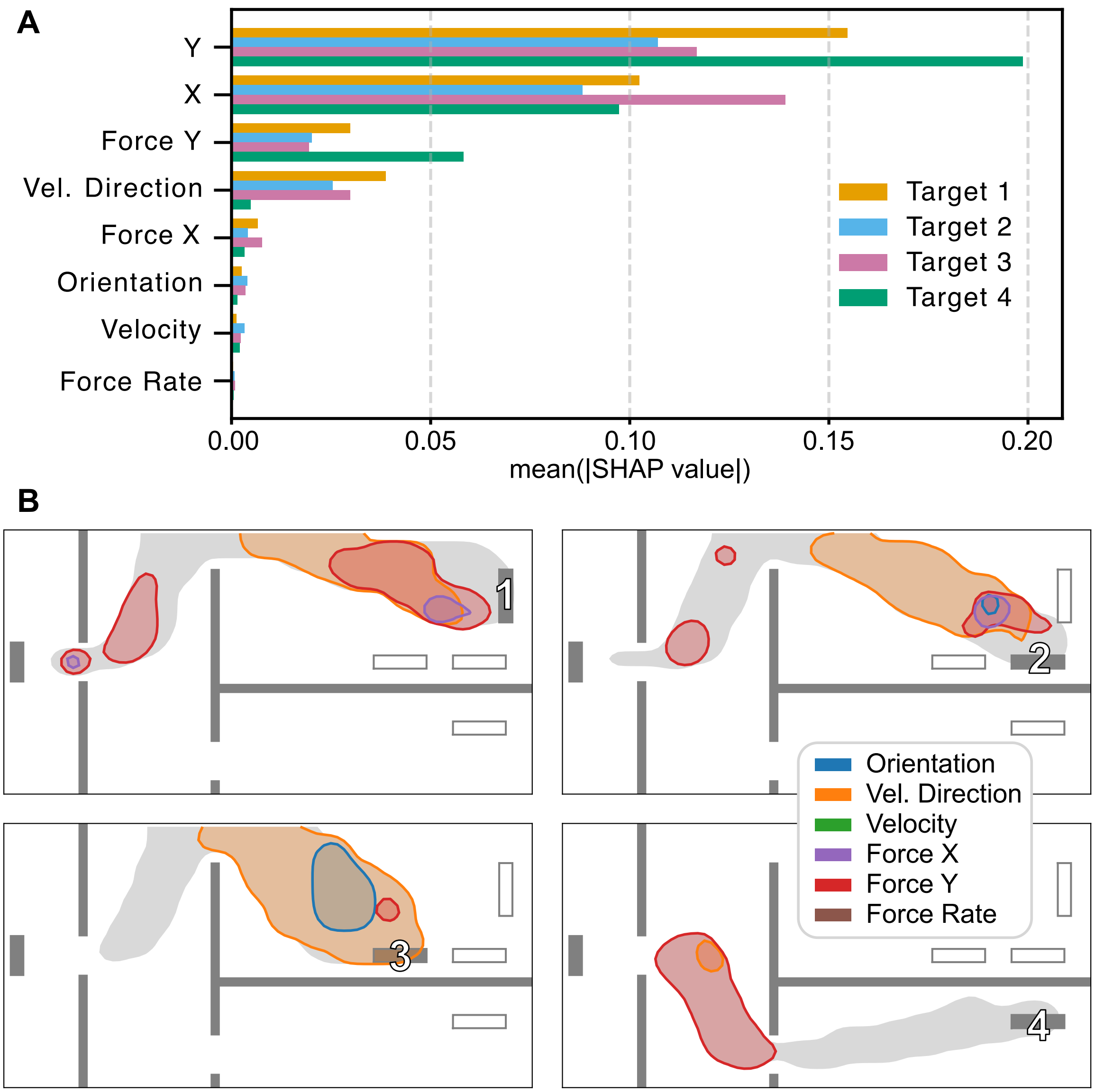


**Figure S16: SHAP analysis explaining the RF predictions and feature contributions in Scene 1. A.** Mean absolute SHAP values for each feature and target, showing the overall importance of each feature. **B.** Spatial regions where each feature substantially increased the model's confidence for each target. Contours indicate regions where the 95th percentile of the signed SHAP value exceeded 0.1 in probability units, color-coded by feature. Table position was excluded to highlight where other features contributed. Gray shading indicates the overall distribution of trajectories.

#### Supplementary Note 5 Distribution of button press positions in Scene 1

In Fig. 6A, the table positions at the time of the button press are shown for Scene 2. Here, we present the corresponding map for Scene 1 (Fig. S17).

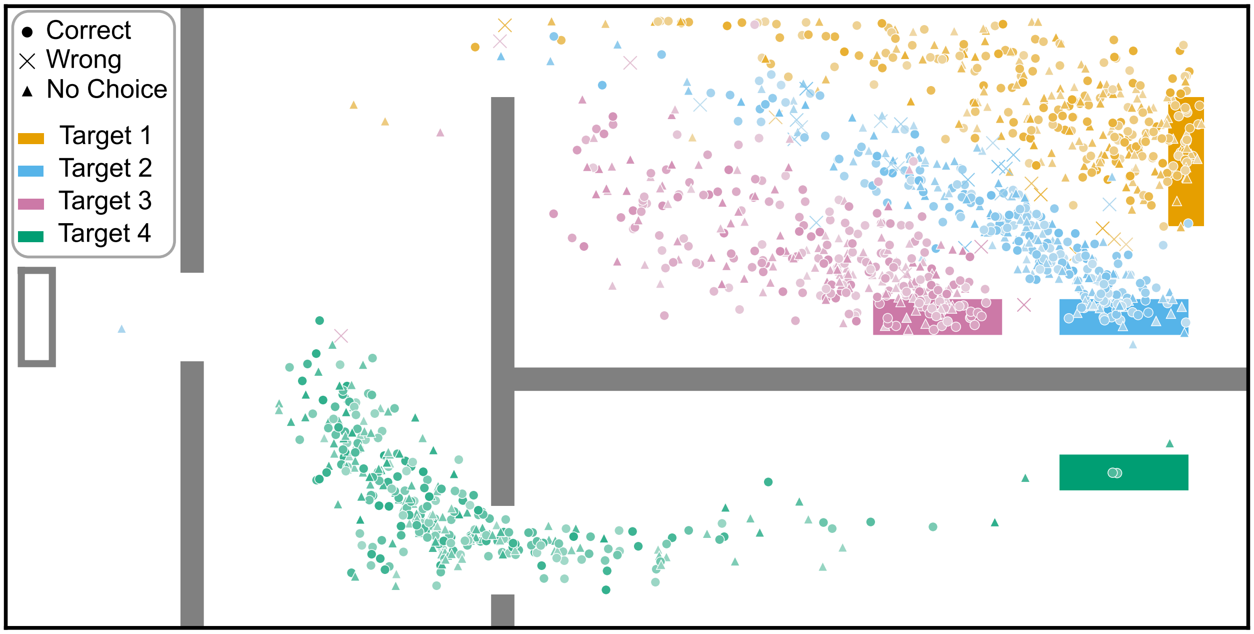


**Figure S17: Table positions at the moment of the button press for Scene 1.** Circles and crosses indicate correct and incorrect choices in the *Button Press-Select* condition, respectively. Triangles indicate *Button Press-Transport* trials, where participants did not report their target choice. Markers are color-coded by target. Lighter and darker shaded markers indicate earlier and later trials, respectively.

### Supplementary Note 6 Dyad-specific signaling performance

A successful signaling strategy should allow the follower to infer the target earlier during the movement. For *Button Press* trials, we quantified this in terms of both time and position, using two metrics: button press time and table $x$-position at button press. In addition, for *Button Press-Select* trials, we quantified target choice accuracy.

In the main text, we reported the Jensen–Shannon (JS) distance between dyad-specific models and the general model, which quantifies how much each dyad's signaling strategy deviated from the general strategy. Here, we ranked dyads according to this JS distance and plotted the corresponding behavioral metrics to examine whether dyad-specific signaling strategies were associated with differences in task performance (Fig. S18).

Overall, target choice accuracy was high across dyads. Button press time and table $x$-position at button press showed greater variability across dyads. Although the relationship was not monotonic, dyads with larger JS distance tended to press the button earlier. For example, D9 showed the shortest button press time, while D1 pressed the button at the earliest table $x$-position, and both dyads had relatively large JS distances from the general model. These results suggest that dyad-specific strategies should not be interpreted as degraded strategies. Instead, distinctive strategies may have enabled more efficient signaling.

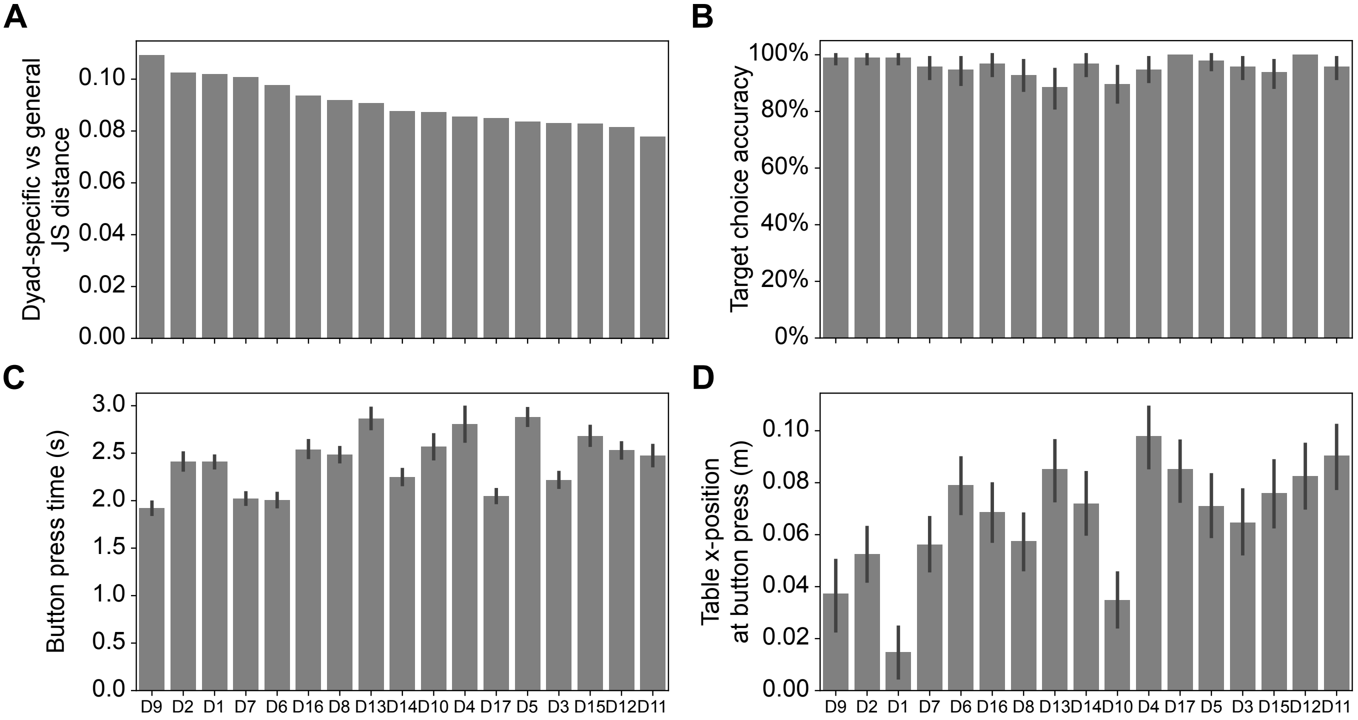


**Figure S18: Dyad-specific signaling performance. A.** JS distance between the dyad-specific model and the general model. Larger JS distance indicates a more distinctive dyad-specific signaling strategy. **B.** Target choice accuracy in *Button Press-Select* trials. **C.** Button press time in *Button Press* trials. **D.** Table $x$-position at button press in *Button Press* trials. All panels are ordered by descending JS distance, from the most to the least unique dyad-specific model.
